# Computational and Molecular Dynamics Simulation Approach To Analyze the Impact of *XPD* Gene Mutation on Protein Stability and Function

**DOI:** 10.1101/2020.07.18.209841

**Authors:** Nagesh Kishan Panchal, Aishwarya Bhale, Vinod Kumar Verma, Syed Sultan Beevi

**Author notes:** **Corresponding Author**, Syed Sultan Beevi PhD, Department of Cancer Biology, KIMS Foundation and Research Centre, KIMS Hospitals, 1-8-31/1, Minister Road, Secunderabad 500 003, Telangana, India. **Co-Corresponding Author**, Vinod Kumar Verma PhD, Department of Cancer Biology, KIMS Foundation and Research Centre, KIMS Hospitals, 1-8-31/1, Minister Road, Secunderabad 500 003, Telangana, India.

## Abstract

*XPD* acts as a functional helicase and aids in unwinding double helix around damaged DNA, leading to efficient DNA repair. Mutations of *XPD* give rise to DNA-repair deficiency diseases and cancer proneness. In this study, cancer-causing missense mutation that could inactivate helicase function and hinder its binding with other complexes were analysed using bioinformatics approach. Rigorous computational methods were employed to understand the molecular pathogenic profile of mutation. The mutant model with the desired mutation was built with I-TASSER. GROMACS 5.0.1 was used to evaluate the effect of a mutation on protein stability and function. Of the 276 missense mutations, 64 were found to be disease-causing. Out of these 64, seven were of cancer-causing mutations. Among these, we evaluated K48R mutation in a computational simulated environment to determine its impact on protein stability and function since K48 position was ascertained to be highly conserved and substitution with arginine could impair the *XPD* activity. Molecular Dynamic Simulation and Essential Dynamics analysis showed that K48R mutation altered protein structural stability and produced conformational drift. Our predictions thus revealed that K48R mutation could impair the *XPD* helicase activity and affect its ability to repair the damaged DNA, thus augmenting the risk for cancer.

## Introduction

The human *XPD* protein, encoded by *ERCC2* is a helicase with 5–3’ polarity [1]. It is a key component of the transcription-coupled nucleotide excision repair (NER) pathway that is critical for DNA damage repair and protects against gene mutations [2]. The crystal structure of *XPD* revealed the presence of two domains (HD1 and HD2) separated by a cleft containing the residues that bind ATP along with 4FeS cluster and Arch domain both of which inserted into the HD1 domain [3]. It is organized within the context of the general transcription factor IIH (TFIIH) and has ATP-dependent helicase activity [4].

*XPD* assumes significance owing to its strikingly different roles during transcription and NER. XPD acts as a fully functional helicase by forming a core component of the transcription factor TFIIH during NER and aids in unwinding the DNA double helix around damaged DNA. Whereas in transcription, it acts as a scaffold for positioning additional factors like CAK complex [5].

Mutations of the human *XPD* gene is often associated with DNA repair deficiency and a high cancer proneness. It is reasonable that minor variations in the DNA repair mechanism owing to mutation may facilitate cancer development. To date almost 100 different mutations have been identified in the *XPD* gene [6,7]. Interestingly, most of the mutations affecting *XPD* are clustered in the C-terminal domain of the protein, which is the pivotal interaction domain of XPD for the p44 subunit of TFIIH [8]. Nonetheless, there is a dearth of literature that could distinguish the most significant mutation(s) responsible for cancer initiation and promotion.

Of late, computational analysis has been emerged as a credible approach to screen for deleterious mutation and ascertain the role of missense mutations on structural changes of a particular protein that would affect its activity. In this study, our objective was to evaluate the potential effects of mutations residing at the different domains of *XPD* that could impair its function and play a role in cancer promotion. We primarily employed multiple computational algorithms such as, Meta-SNP, Pmut, and Provean to categorize deleterious mutations and later used FATHMM server to identify cancer-promoting/cancer-driving mutations among disease-causing mutations. Furthermore, we used mCSM, SDM2, STRUM, and Mutation Assessor to predict the impact of the mutation on protein structure and stability. A mutant model with desired missense mutation was built with I-TASSER server and a 3D mutated structure was simulated using GROMACS 5.0.1 to investigate the mechanism of structural consequences of mutation on XPD protein stability and function.

## Materials and Methods

### Collection of dataset

The mutations and associated information of *XPD* missense mutations were collected from HGMD (www.hgmd.cf.ac.uk/ac/index.php), COSMIC (https://cancer.sanger.ac.uk/cosmic), PubMed (https://pubmed.ncbi.nlm.nih.gov/). Sequence information in FASTA format was retrieved from UniProtKB (UniPort ID: P18074) (https://www.uniprot.org/) whereas structural information of *XPD* was obtained from Protein Data Bank (PDB ID: 5ivw) (http://www.rcsb.org/).

### Computational pathogenicity analysis of mutations

Computational methods to understand the impact of mutations on proteins are important for classifying and prioritizing pathogenic and neutral single-nucleotide mutations.

Meta-SNP (http://snps.biofold.org/meta-snp/) incorporates the best-performing prediction algorithms to classify the deleteriousness of mutations in a protein. In addition to its algorithm, this method integrates a variety of other algorithms, including SNAP Prediction, PhD-SNP Prediction, PANTHER Prediction, and SIFT Prediction. Meta-SNP is a 100-tree Random Forest WEKA library operation, trained on SV-2009 using 20-fold cross-validation. The predictor outputs the probability that a given mutation is disease-related, where scores >0.5 specify that the given mutation is disease-causing [9].

PMut (http://mmb.irbbarcelona.org/PMut) is a neural network-based program for the annotation of pathological mutations on proteins. It employs three parameters such as a mutation descriptors, solvent accessibility, and residue/sequence properties to estimate the pathogenicity indexes of given input mutation data with a prediction score between 0 and 1; mutations scoring from 0 to 0.5 are classified as neutral and those scoring from 0.5 to 1 are classified as pathological [10].

PROVEAN (Protein variation effect analyser - http://provean.jcvi.org/seq_submit.php) is a web server that uses an alignment-based score approach to predict whether an amino acid substitution or indel has an impact on the biological function of a protein [11]. Mutations are predicted as deleterious when the final scale is below the threshold value of −2.5.

Mutation Assessor (http://mutationassessor.org/r3/) calculates the functional impact of amino acid substitutions in proteins. It is based on evolutionary conservation of the affected amino acid in protein homologs, specifying a rough estimate of a probability that the mutation has a phenotypic consequence at the level of an organism. It utilizes information based on the analysis of evolutionary conservation patterns in protein family multiple-sequence alignments. The analysis results in a functional impact score based on evolutionary information (FIS), variant conservation score (VC), and variant specificity score that classifies the mutations as neutral, low, medium or high [12].

### Protein stability analysis

Studying the effects of mutation on protein stability and function is important in understanding its role in disease.

SDM2 (Site Directed Mutator -http://marid.bioc.cam.ac.uk/sdm2) is a computational method that analyses the variation of amino acid substitution occurring at a specific structural environment that is tolerated within the family of homologous proteins of known 3D structures and convert them into substitution probability tables which are used a quantitative measure for predicting the protein stability upon mutation [13].

mCSM (http://biosig.unimelb.edu.au/mcsm/) is a computational approach to predict the impact of missense mutation on protein stability and interactions between associating partners. This relies on graph-based signatures and encode distance patterns between atoms and is used to represent the protein residue environment and to train predictive models. mCSM evaluates the impact not only on protein stability but also on protein-protein and protein-nucleic acid interactions to understand the role of mutations in disease [14].

STRUM (https://zhanglab.ccmb.med.umich.edu/STRUM/) is a computational method for predicting stability changes (ΔΔG) of a protein upon single-point mutation. It takes on a gradient boosting regression approach using a variety of features at different levels of evolutionary information and structural resolution [15].

DUET (http://biosig.unimelb.edu.au/duet/), is an integrated computational approach for predicting the effects of missense mutations on protein stability. DUET combines mCSM and SDM in a consensus prediction, by consolidating the results of the separate methods in an optimized predictor using Support Vector Machines (SVMs) trained with Sequential Minimal Optimization [16].

### Cancer association analysis

FATHMM server (http://fathmm.biocompute.org.uk/cancer.html) was used to check the cancer association of the deleterious mutations. The server returns predictions capable of discriminating between cancer-promoting/cancer-driving mutation and other germline polymorphism. [17].

### Analysis of sequence conservation

The ConSurf web server (http://consurf.tau.ac.il) analyses the evolutionary pattern of the amino acids of the protein to reveal regions that are important for structure and/or function. Starting from a query sequence or structure, the server automatically collects homologs, infers their multiple sequence alignment, and reconstructs a phylogenetic tree that reflects their evolutionary relations. These data are then used, within a probabilistic framework, to estimate the evolutionary rates of each sequence position [18].

### Modeling mutation location on protein structure

The crystal structure of native *XPD* was downloaded from Protein data bank and was fixed using WHATIF server. The mutant model with desired mutation (K48R) was built using I-TASSER server [19] which uses multiple threading approach LOMETS with full-length atomic models constructed by iterative template-based fragment assembly simulations.

### Quality assessment of Protein Model

A quality check of the mutant protein model which was obtained from I-TASSER server was performed using the Rampage server.

### Molecular Dynamic (MD) Simulation

The best-scored mutant model structure obtained from I-TASSER was chosen as the starting coordinates for further analysis. To investigate the mechanism of structural consequences of the mutations on *XPD*, MD simulation of the K48R mutation was performed using GROMACS 5.0.1 [20] with OPLS force field where box type was cubic and size of the box was 2.5 – nm. The SPC water model was used and the system was neutralized by substituting a solvent molecule from Cl- ions. A periodic boundary condition was applied in all directions. Subsequently a maximum of 50,000 steps of energy minimization was carried out for the predicted models using a conjugate gradient algorithm followed by steepest descent minimization. Then equilibration was carried out for 1000 ps for each system with NVT (constant number of particles, volume, and temperature) and the temperature was kept constant using a Berendsen thermostat followed by NPT (constant number of particles, pressure, and temperature) with Parinello-Rahman pressure coupling. Finally the equilibrated system was subjected to 25 ns MD simulation with time –step of 2 fs.

### Analysis of MD trajectories

Comparative analysis of the Structural deviation in native and mutant (K48R) structure such as root mean square deviation (RMSD), root mean square fluctuation (RMSF), radius of gyration (Rg), solvent accessible surface area (SASA), Hydrogen bond, etc., were computed using GROMACS associated utility packages such as g_rms, g_rmsf, g_gyrate, g_sas, and g_hbond. GRACE software was used for plotting all graphs (http://plasma-gate.weizmann.ac.il/Grace/).

### Secondary structure analysis

The software tool DSSP (Database of Secondary Structure in Proteins) by Kabsch and Sander was applied to analyse the change in secondary structure patterns in the native and mutant protein. DSSP tool employs H-bonding patterns and various other geometrical features to assign secondary structure labels to the residues of a protein. Secondary structure pattern was plotted between native and mutant and superimposed at the beginning of the simulation and at specific time points where the conformational drifts occurred at a higher range.

### Principal Component analysis

The principal component analysis is an unsupervised statistical technique for finding patterns in high-dimensional data. It is one of the effective methods for finding global and correlation motion by obtaining standard information from the MD trajectories through dimensionality reduction method. It has been used to distinguish the large scale collection motions from the random thermal fluctuations.

PCA method is based on the calculation and diagonalization of the covariance matrix. The elements Cij in this matrix are defined by.

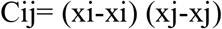

Where xi (xj) is the coordinate of the ith (jth) atom and represents an ensemble average. In general, the few initial principal components (PCs) are responsible for the most important conformational changes of the bimolecular system by storing eigenvectors with the highest eigenvalues. g_covar is GROMACS utility package that was used for generating the covariance matrix using the protein backbone as a reference structure for the rotational fit and g_anaeig was used to analyse and plot the eigenvectors.

## Results

Our results illustrate the impact of missense mutations in the *XPD* gene on its protein structure, stability, and function. Deleterious or pathogenic spectrum, protein stability, and cancer-causing mutations were screened using advanced computational methods.

### Dataset retrieval of missense mutations

We retrieved a list of 276 mutations from the public database of the *XPD* gene located at different coding regions for our investigation.

### Analysis of disease-causing mutations

The pathogenic impact of missense mutations can be determined by evaluating the significance of the amino acids they affect. Summary of 276 missense mutations that were analysed by Meta-SNP (PATHER, PhD-SNP, SIFT, SNAP), Pmut, and Provean is represented as graph [Figure 1A] and detail dataset is presented as [Table1]. Results obtained from these servers showed that 67 mutations were predicted to be deleterious or disease-causing and were residing on different domains of *XPD* as illustrated in [Figure 1B].

**Figure 1.**
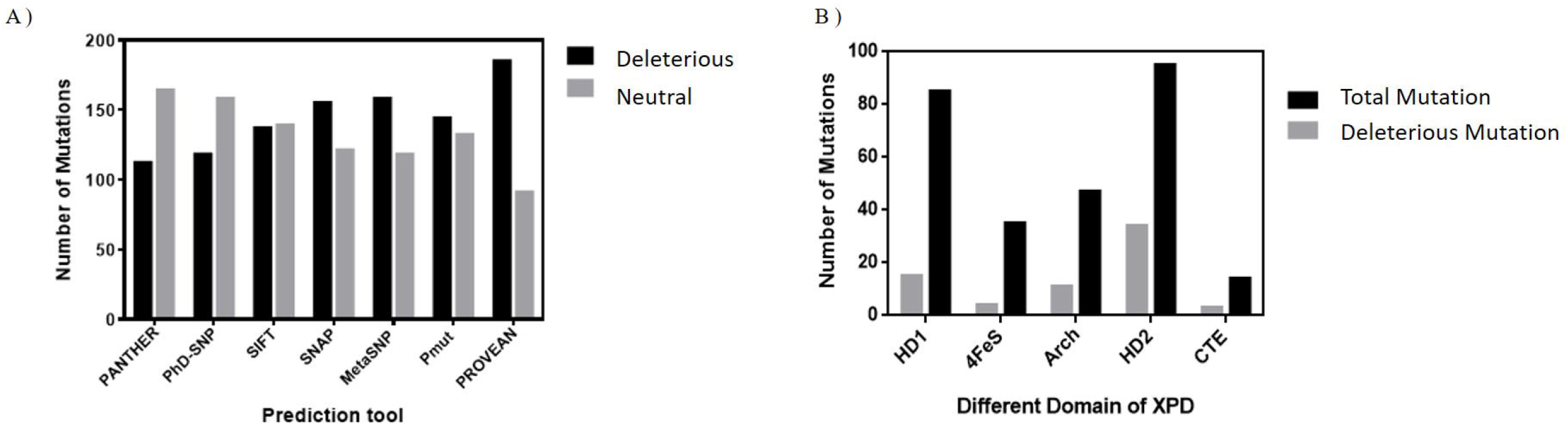
Computational prediction and screening of mutation in XPD **A:** Screening of deleterious & neutral mutations using PATHER, PHD-SNP, SIFT, SNAP, Pmut, PROVEAN **B:** Deleterious mutations residing on different domains of XPD

**Table 1.** Prediction of Deleterious Mutation using computational tools

| Prediction of Deleterious Mutation |  |  |  |  |  |  |  |  |  |  |  |  |  |  |  |
| --- | --- | --- | --- | --- | --- | --- | --- | --- | --- | --- | --- | --- | --- | --- | --- |
| Sr.no | Mutants | Meta-SNP |  |  |  |  |  |  |  |  |  | Pmut |  | Provean |  |
|  |  | PANTHER |  | PhD-SNP |  | SIFT |  | SNAP |  | Meta-SNP |  |  |  |  |  |
|  |  | Prediction | Score | Prediction | Score | Prediction | Score | Prediction | Score | Prediction | Score | Prediction | Score | Prediction | Score |
| 1 | E20K | Neutral | 0.427 | Disease | 0.821 | Disease | 0 | Disease | 0.655 | Disease | 0.756 | Disease | 0.64 | Deleterious | -3.107 |
| 2 | Q21E | Disease | 0.978 | Disease | 0.84 | Disease | 0 | Disease | 0.71 | Disease | 0.799 | Disease | 0.78 | Neutral | -2.33 |
| 3 | S23F | Neutral | 0.458 | Disease | 0.533 | Disease | 0.04 | Neutral | 0.485 | Disease | 0.51 | Disease | 0.54 | Neutral | -2.229 |
| 4 | M25L | Neutral | 0.373 | Disease | 0.855 | Disease | 0.04 | Neutral | 0.525 | Disease | 0.572 | Disease | 0.52 | Deleterious | -2.578 |
| 5 | R26Q | Neutral | 0.11 | Neutral | 0.23 | Neutral | 0.6 | Neutral | 0.125 | Neutral | 0.353 | Neutral | 0.15 | Neutral | 0.21 |
| 6 | R26W | Neutral | 0.446 | Disease | 0.534 | Disease | 0.02 | Neutral | 0.125 | Neutral | 0.353 | Neutral | 0.5 | Neutral | -0.691 |
| 7 | R30C | Disease | 0.769 | Disease | 0.679 | Disease | 0 | Disease | 0.575 | Disease | 0.722 | Disease | 0.77 | Deleterious | -5.371 |
| 8 | T31M | Disease | 0.709 | Disease | 0.596 | Disease | 0.01 | Disease | 0.565 | Disease | 0.707 | Disease | 0.83 | Neutral | -2.37 |
| 9 | H37R | Neutral | 0.263 | Disease | 0.681 | Disease | 0 | Disease | 0.625 | Disease | 0.721 | Disease | 0.76 | Deleterious | -6.515 |
| 10 | V39I | Neutral | 0.145 | Neutral | 0.084 | Neutral | 0.36 | Neutral | 0.215 | Neutral | 0.192 | Neutral | 0.28 | Neutral | 0.025 |
| 11 | G47R | Disease | 0.893 | Disease | 0.893 | Disease | 0 | Disease | 0.855 | Disease | 0.878 | Disease | 0.9 | Deleterious | -7.219 |
| 12 | K48R | Disease | 0.511 | Disease | 0.7 | Disease | 0 | Disease | 0.81 | Disease | 0.774 | Disease | 0.9 | Deleterious | -2.707 |
| 13 | T49I | Disease | 0.762 | Disease | 0.825 | Disease | 0.04 | Disease | 0.735 | Disease | 0.774 | Disease | 0.9 | Deleterious | -5.414 |
| 14 | L55P | Neutral | 0.261 | Disease | 0.894 | Disease | 0.01 | Disease | 0.55 | Disease | 0.606 | Disease | 0.78 | Deleterious | -6.317 |
| 15 | M57K | Neutral | 0.166 | Disease | 0.628 | Disease | 0.01 | Disease | 0.55 | Disease | 0.649 | Neutral | 0.26 | Deleterious | -2.642 |
| 16 | P64L | Neutral | 0.446 | Disease | 0.863 | Disease | 0 | Disease | 0.57 | Disease | 0.739 | Disease | 0.68 | Deleterious | -8.908 |
| 17 | E66V | Neutral | 0.101 | Disease | 0.726 | Neutral | 0.12 | Disease | 0.51 | Disease | 0.585 | Disease | 0.5 | Neutral | -2.197 |
| 18 | R75I | Disease | 0.715 | Disease | 0.881 | Disease | 0 | Disease | 0.795 | Disease | 0.841 | Disease | 0.87 | Deleterious | -7.303 |
| 19 | T76A | Neutral | 0.361 | Disease | 0.81 | Disease | 0.01 | Disease | 0.8 | Disease | 0.789 | Disease | 0.88 | Deleterious | -4.564 |
| 20 | P78S | Neutral | 0.102 | Neutral | 0.105 | Neutral | 0.42 | Neutral | 0.34 | Neutral | 0.263 | Neutral | 0.29 | Deleterious | -5.944 |
| 21 | E86Q | Disease | 0.772 | Disease | 0.806 | Disease | 0 | Disease | 0.64 | Disease | 0.576 | Neutral | 0.38 | Deleterious | -2.755 |
| 22 | R88G | Disease | 0.594 | Disease | 0.695 | Neutral | 0.07 | Disease | 0.54 | Disease | 0.657 | Disease | 0.59 | Deleterious | -4.462 |
| 23 | N92S | Neutral | 0.114 | Neutral | 0.103 | Neutral | 0.3 | Neutral | 0.19 | Neutral | 0.174 | Neutral | 0.25 | Neutral | -0.181 |
| 24 | F93L | Neutral | 0.194 | Neutral | 0.441 | Neutral | 0.13 | Neutral | 0.47 | Neutral | 0.425 | Neutral | 0.35 | Deleterious | -3.083 |
| 25 | Y94S | Neutral | 0.166 | Neutral | 0.487 | Neutral | 0.36 | Disease | 0.55 | Neutral | 0.444 | Disease | 0.59 | Deleterious | -4.081 |
| 26 | E95D | Neutral | 0.243 | Neutral | 0.334 | Neutral | 0.38 | Neutral | 0.255 | Neutral | 0.419 | Neutral | 0.29 | Neutral | -1.63 |
| 27 | Q97L | Neutral | 0.31 | Neutral | 0.253 | Neutral | 0.28 | Neutral | 0.38 | Neutral | 0.374 | Disease | 0.52 | Deleterious | -4.035 |
| 28 | E98D | Neutral | 0.28 | Neutral | 0.253 | Neutral | 0.28 | Neutral | 0.38 | Neutral | 0.374 | Neutral | 0.14 | Neutral | -0.321 |
| 29 | K101T | Neutral | 0.115 | Neutral | 0.331 | Neutral | 0.49 | Neutral | 0.265 | Neutral | 0.408 | Neutral | 0.34 | Neutral | -0.878 |
| 30 | P103L | Neutral | 0.28 | Neutral | 0.259 | Neutral | 0.23 | Neutral | 0.41 | Neutral | 0.407 | Neutral | 0.49 | Neutral | -1.998 |
| 31 | P103R | Neutral | 0.198 | Neutral | 0.278 | Neutral | 0.46 | Neutral | 0.385 | Neutral | 0.403 | Neutral | 0.34 | Neutral | 0.252 |
| 32 | R112C | Disease | 0.887 | Disease | 0.89 | Disease | 0 | Disease | 0.835 | Disease | 0.885 | Disease | 0.94 | Deleterious | -7.231 |
| 33 | R112G | Disease | 0.691 | Disease | 0.844 | Disease | 0.04 | Disease | 0.79 | Disease | 0.82 | Disease | 0.92 | Deleterious | -6.329 |
| 34 | R112H | Disease | 0.741 | Disease | 0.88 | Disease | 0 | Disease | 0.81 | Disease | 0.826 | Disease | 0.87 | Deleterious | -4.526 |
| 35 | H118R | Neutral | 0.314 | Disease | 0.7 | Disease | 0 | Disease | 0.77 | Disease | 0.654 | Disease | 0.83 | Deleterious | -5.333 |
| 36 | P123T | Neutral | 0.107 | Neutral | 0.143 | Neutral | 0.47 | Neutral | 0.22 | Neutral | 0.199 | Neutral | 0.16 | Neutral | 0.119 |
| 37 | R125H | Neutral | 0.206 | Neutral | 0.406 | Disease | 0.04 | Disease | 0.625 | Neutral | 0.452 | Neutral | 0.46 | Deleterious | -3.104 |
| 38 | D131N | Disease | 0.504 | Neutral | 0.215 | Disease | 0 | Disease | 0.7 | Neutral | 0.315 | Neutral | 0.42 | Deleterious | -4.576 |
| 39 | C134T | Disease | 0.611 | Disease | 0.942 | Disease | 0.03 | Disease | 0.815 | Disease | 0.866 | Disease | 0.82 | Deleterious | -9.151 |
| 40 | H135D | Neutral | 0.219 | Disease | 0.667 | Neutral | 0.06 | Disease | 0.565 | Disease | 0.519 | Disease | 0.62 | Deleterious | -3.182 |
| 41 | S136N | Neutral | 0.228 | Neutral | 0.309 | Neutral | 0.24 | Neutral | 0.395 | Neutral | 0.422 | Neutral | 0.38 | Neutral | -0.321 |
| 42 | Y141C | Disease | 0.835 | Disease | 0.536 | Neutral | 0.17 | Disease | 0.525 | Disease | 0.63 | Disease | 0.61 | Deleterious | -4.649 |
| 43 | V142L | Neutral | 0.26 | Neutral | 0.296 | Neutral | 0.71 | Neutral | 0.19 | Neutral | 0.412 | Neutral | 0.34 | Neutral | -2.134 |
| 44 | R143Q | Neutral | 0.462 | Disease | 0.634 | Neutral | 0.53 | Neutral | 0.355 | Neutral | 0.496 | Neutral | 0.45 | Deleterious | -3.193 |
| 45 | R143G | Disease | 0.705 | Disease | 0.736 | Neutral | 0.26 | Disease | 0.565 | Disease | 0.619 | Neutral | 0.39 | Deleterious | -5.56 |
| 46 | R143T | Disease | 0.625 | Disease | 0.674 | Neutral | 0.43 | Neutral | 0.425 | Disease | 0.558 | Disease | 0.4 | Deleterious | -4.592 |
| 47 | A144V | Neutral | 0.275 | Neutral | 0.292 | Neutral | 0.28 | Neutral | 0.28 | Neutral | 0.405 | Neutral | 0.33 | Neutral | -1.552 |
| 48 | H148T | Neutral | 0.11 | Neutral | 0.319 | Neutral | 0.61 | Neutral | 0.31 | Neutral | 0.405 | Neutral | 0.09 | Neutral | -0.419 |
| 49 | D149N | Neutral | 0.244 | Neutral | 0.388 | Neutral | 0.6 | Neutral | 0.32 | Neutral | 0.421 | Neutral | 0.27 | Neutral | -1.454 |
| 51 | L152R | Neutral | 0.47 | Disease | 0.694 | Neutral | 0.45 | Neutral | 0.335 | Disease | 0.538 | Disease | 0.65 | Deleterious | -2.984 |
| 52 | Y158C | Disease | 0.87 | Disease | 0.765 | Neutral | 0.17 | Disease | 0.555 | Disease | 0.734 | Disease | 0.55 | Deleterious | -5.633 |
| 53 | H164R | Neutral | 0.11 | Neutral | 0.224 | Neutral | 0.37 | Neutral | 0.47 | Neutral | 0.362 | Neutral | 0.2 | Neutral | -1.506 |
| 54 | E167Q | Neutral | 0.181 | Neutral | 0.293 | Neutral | 0.58 | Neutral | 0.225 | Neutral | 0.413 | Neutral | 0.25 | Neutral | -1.383 |
| 55 | V168M | Neutral | 0.109 | Neutral | 0.154 | Neutral | 0.12 | Neutral | 0.23 | Neutral | 0.189 | Neutral | 0.24 | Neutral | 0.321 |
| 56 | L170F | Disease | 0.566 | Neutral | 0.463 | Neutral | 0.67 | Neutral | 0.22 | Disease | 0.574 | Neutral | 0.38 | Deleterious | -2.731 |
| 57 | L170R | Disease | 0.857 | Disease | 0.879 | Neutral | 0.21 | Disease | 0.555 | Disease | 0.765 | Neutral | 0.48 | Deleterious | -4.474 |
| 58 | A172T | Neutral | 0.166 | Neutral | 0.211 | Neutral | 0.49 | Neutral | 0.3 | Neutral | 0.311 | Neutral | 0.23 | Neutral | 0.034 |
| 59 | N176S | Neutral | 0.202 | Neutral | 0.189 | Neutral | 0.86 | Neutral | 0.285 | Neutral | 0.283 | Neutral | 0.25 | Neutral | -0.756 |
| 60 | D178E | Neutral | 0.215 | Neutral | 0.216 | Neutral | 0.8 | Neutral | 0.25 | Neutral | 0.252 | Neutral | 0.3 | Neutral | -1.006 |
| 60 | G184V | Neutral | 0.403 | Disease | 0.92 | Disease | 0 | Disease | 0.77 | Disease | 0.787 | Disease | 0.94 | Deleterious | -6.724 |
| 61 | R185T | Neutral | 0.256 | Neutral | 0.465 | Neutral | 0.2 | Disease | 0.56 | Neutral | 0.456 | Disease | 0.65 | Deleterious | -2.567 |
| 62 | R186C | Disease | 0.611 | Disease | 0.627 | Disease | 0.02 | Disease | 0.655 | Disease | 0.696 | Disease | 0.7 | Deleterious | -3.395 |
| 63 | R186H | Neutral | 0.28 | Neutral | 0.347 | Disease | 0.05 | Disease | 0.58 | Neutral | 0.446 | Neutral | 0.39 | Neutral | -1.529 |
| 64 | Q187R | Neutral | 0.184 | Neutral | 0.255 | Neutral | 0.42 | Neutral | 0.355 | Neutral | 0.36 | Neutral | 0.29 | Neutral | 0.539 |
| 65 | S198L | Neutral | 0.132 | Neutral | 0.435 | Neutral | 0.71 | Neutral | 0.22 | Neutral | 0.424 | Neutral | 0.49 | Neutral | -0.027 |
| 66 | I199M | Neutral | 0.307 | Neutral | 0.351 | Neutral | 0.3 | Neutral | 0.43 | Neutral | 0.428 | Neutral | 0.35 | Neutral | -1.543 |
| 67 | H201T | Neutral | 0.199 | Neutral | 0.402 | Neutral | 0.39 | Disease | 0.665 | Neutral | 0.437 | Neutral | 0.47 | Deleterious | -4.156 |
| 68 | A202V | Disease | 0.517 | Disease | 0.883 | Neutral | 0.12 | Neutral | 0.49 | Disease | 0.594 | Disease | 0.68 | Deleterious | -2.761 |
| 69 | N203S | Neutral | 0.276 | Disease | 0.713 | Disease | 0.05 | Disease | 0.615 | Disease | 0.724 | Disease | 0.66 | Deleterious | -2.959 |
| 70 | Y209C | Disease | 0.923 | Disease | 0.952 | Disease | 0 | Disease | 0.84 | Disease | 0.901 | Disease | 0.94 | Deleterious | -7.981 |
| 71 | P215S | Neutral | 0.481 | Disease | 0.715 | Disease | 0 | Disease | 0.62 | Disease | 0.583 | Disease | 0.43 | Deleterious | -7.447 |
| 72 | A218V | Neutral | 0.434 | Disease | 0.72 | Disease | 0.03 | Disease | 0.605 | Disease | 0.719 | Disease | 0.84 | Deleterious | -3.665 |
| 73 | L220M | Neutral | 0.329 | Neutral | 0.179 | Neutral | 0.21 | Neutral | 0.225 | Neutral | 0.271 | Neutral | 0.26 | Neutral | -0.387 |
| 74 | L220V | Neutral | 0.132 | Neutral | 0.177 | Neutral | 0.7 | Neutral | 0.185 | Neutral | 0.227 | Neutral | 0.26 | Neutral | -0.786 |
| 75 | S222A | Neutral | 0.352 | Neutral | 0.295 | Disease | 0.02 | Disease | 0.6 | Neutral | 0.154 | Neutral | 0.49 | Deleterious | -2.793 |
| 76 | A226T | NA | NA | Neutral | 0.193 | Neutral | 0.52 | Neutral | 0.33 | Neutral | 0.154 | Neutral | 0.38 | Neutral | 0.078 |
| 77 | R227C | Disease | 0.773 | Disease | 0.587 | Disease | 0.01 | Disease | 0.595 | Disease | 0.726 | Disease | 0.78 | Deleterious | -5.212 |
| 78 | R227H | Neutral | 0.445 | Neutral | 0.359 | Disease | 0.02 | Disease | 0.525 | Neutral | 0.456 | Neutral | 0.35 | Deleterious | -3.164 |
| 79 | K228T | Neutral | 0.292 | Neutral | 0.256 | Neutral | 0.2 | Neutral | 0.455 | Neutral | 0.414 | Neutral | 0.41 | Neutral | -1.235 |
| 80 | A229V | Neutral | 0.174 | Neutral | 0.361 | Disease | 0.03 | Disease | 0.6 | Neutral | 0.452 | Disease | 0.58 | Deleterious | -2.626 |
| 81 | V231M | Disease | 0.611 | Disease | 0.713 | Disease | 0 | Disease | 0.7 | Disease | 0.726 | Disease | 0.84 | Deleterious | -2.759 |
| 82 | N241I | Disease | 0.699 | Disease | 0.761 | Neutral | 0.08 | Neutral | 0.38 | Disease | 0.617 | Disease | 0.79 | Deleterious | -8.528 |
| 83 | I244M | Neutral | 0.487 | Neutral | 0.404 | Neutral | 0.18 | Neutral | 0.285 | Neutral | 0.43 | Disease | 0.62 | Deleterious | -2.701 |
| 84 | I244T | Neutral | 0.471 | Disease | 0.612 | Disease | 0.03 | Disease | 0.53 | Neutral | 0.495 | Disease | 0.63 | Deleterious | -4.296 |
| 85 | D245G | Neutral | 0.414 | Disease | 0.792 | Disease | 0.05 | Disease | 0.575 | Disease | 0.73 | Disease | 0.68 | Deleterious | -4.383 |
| 86 | M247T | Neutral | 0.312 | Neutral | 0.429 | Disease | 0.02 | Disease | 0.655 | Neutral | 0.472 | Neutral | 0.45 | Deleterious | -4.632 |
| 87 | N250S | Neutral | 0.382 | Neutral | 0.251 | Neutral | 0.47 | Neutral | 0.345 | Neutral | 0.394 | Neutral | 0.29 | Neutral | -0.376 |
| 88 | T252P | Neutral | 0.419 | Disease | 0.8 | Neutral | 0.19 | Neutral | 0.44 | Disease | 0.584 | Neutral | 0.48 | Deleterious | -2.883 |
| 89 | R253C | Disease | 0.784 | Disease | 0.629 | Disease | 0.01 | Disease | 0.655 | Disease | 0.72 | Disease | 0.67 | Deleterious | -5.828 |
| 90 | R254G | Neutral | 0.312 | Neutral | 0.438 | Neutral | 0.37 | Disease | 0.565 | Neutral | 0.47 | Neutral | 0.43 | Deleterious | -4.004 |
| 91 | R258T | Neutral | 0.309 | Neutral | 0.336 | Neutral | 0.3 | Neutral | 0.49 | Neutral | 0.423 | Disease | 0.57 | Neutral | -1.926 |
| 92 | C259T | Neutral | 0.2 | Disease | 0.549 | Disease | 0.01 | Disease | 0.67 | Disease | 0.657 | Disease | 0.66 | Neutral | -1.831 |
| 93 | K274E | Neutral | 0.184 | Disease | 0.57 | Disease | 0.4 | Disease | 0.62 | Neutral | 0.469 | Neutral | 0.7 | Deleterious | -3.464 |
| 94 | Q279P | Neutral | 0.299 | Disease | 0.687 | Neutral | 0.13 | Disease | 0.585 | Disease | 0.566 | Neutral | 0.33 | Neutral | -2.453 |
| 95 | R282Q | Neutral | 0.103 | Neutral | 0.166 | Neutral | 1 | Neutral | 0.2 | Neutral | 0.177 | Neutral | 0.24 | Neutral | 0.139 |
| 96 | D283V | Neutral | 0.302 | Neutral | 0.431 | Neutral | 0.13 | Neutral | 0.42 | Neutral | 0.465 | Disease | 0.52 | Deleterious | -3.85 |
| 97 | R286T | Neutral | 0.168 | Neutral | 0.29 | Disease | 0 | Disease | 0.635 | Disease | 0.726 | Neutral | 0.38 | Neutral | -0.676 |
| 98 | R287Y | Disease | 0.644 | Disease | 0.602 | Disease | 0 | Disease | 0.635 | Disease | 0.726 | Disease | 0.65 | Deleterious | -5.585 |
| 99 | A297V | Neutral | 0.104 | Neutral | 0.09 | Neutral | 0.29 | Neutral | 0.17 | Neutral | 0.149 | Neutral | 0.24 | Neutral | -0.037 |
| 100 | R299T | Neutral | 0.191 | Neutral | 0.455 | Neutral | 0.17 | Disease | 0.56 | Neutral | 0.428 | Neutral | 0.44 | Deleterious | -3.69 |
| 101 | E300Q | Neutral | 0.153 | Neutral | 0.307 | Neutral | 0.58 | Neutral | 0.24 | Neutral | 0.395 | Neutral | 0.32 | Neutral | -1.652 |
| 102 | T301K | Neutral | 0.154 | Neutral | 0.527 | Neutral | 0.79 | Neutral | 0.41 | Neutral | 0.433 | Neutral | 0.34 | Deleterious | -2.928 |
| 103 | T301M | Neutral | 0.476 | Neutral | 0.323 | Neutral | 0.09 | Neutral | 0.455 | Neutral | 0.402 | Neutral | 0.46 | Deleterious | -3.565 |
| 104 | N307K | Neutral | 0.38 | Disease | 0.543 | Neutral | 0.05 | Disease | 0.565 | Neutral | 0.484 | Disease | 0.59 | Deleterious | -5.093 |
| 105 | D312Y | Neutral | 0.499 | Disease | 0.65 | Disease | 0.01 | Disease | 0.73 | Disease | 0.731 | Disease | 0.55 | Deleterious | -6.24 |
| 106 | V314A | Disease | 0.525 | Neutral | 0.227 | Neutral | 0.13 | Disease | 0.515 | Neutral | 0.155 | Neutral | 0.31 | Deleterious | -3.413 |
| 107 | Q316E | Neutral | 0.184 | Neutral | 0.168 | Neutral | 0.36 | Neutral | 0.27 | Neutral | 0.173 | Neutral | 0.23 | Neutral | -0.63 |
| 108 | Q316K | Neutral | 0.193 | Neutral | 0.144 | Neutral | 1 | Neutral | 0.205 | Neutral | 0.155 | Neutral | 0.25 | Neutral | -0.369 |
| 109 | I323M | Disease | 0.616 | Disease | 0.586 | Disease | 0 | Disease | 0.72 | Disease | 0.617 | Disease | 0.42 | Neutral | -2.307 |
| 110 | R324H | Disease | 0.792 | Disease | 0.762 | Disease | 0.01 | Disease | 0.655 | Disease | 0.688 | Disease | 0.53 | Deleterious | -4.51 |
| 111 | T325M | Neutral | 0.375 | Neutral | 0.295 | Disease | 0.02 | Disease | 0.505 | Neutral | 0.429 | Neutral | 0.36 | Neutral | -1.12 |
| 112 | E327G | Neutral | 0.46 | Disease | 0.755 | Neutral | 0.05 | Disease | 0.615 | Disease | 0.603 | Disease | 0.56 | Deleterious | -5.175 |
| 113 | R335W | Disease | 0.84 | Disease | 0.924 | Neutral | 0.11 | Neutral | 0.305 | Disease | 0.511 | Neutral | 0.45 | Deleterious | -7.217 |
| 114 | R345H | Disease | 0.524 | Disease | 0.618 | Neutral | 0.11 | Neutral | 0.305 | Disease | 0.751 | Disease | 0.42 | Deleterious | -3.573 |
| 115 | F394L | Disease | 0.509 | Disease | 0.59 | Neutral | 0.31 | Neutral | 0.4 | Disease | 0.503 | Disease | 0.65 | Deleterious | -4.418 |
| 116 | P396L | Disease | 0.526 | Disease | 0.668 | Neutral | 0.82 | Neutral | 0.41 | Disease | 0.541 | Disease | 0.55 | Deleterious | -3.28 |
| 117 | P396S | Neutral | 0.115 | Neutral | 0.4 | Neutral | 0.72 | Neutral | 0.22 | Neutral | 0.414 | Neutral | 0.32 | Neutral | -0.618 |
| 118 | T398I | Neutral | 0.372 | Neutral | 0.411 | Neutral | 0.29 | Neutral | 0.205 | Neutral | 0.451 | Neutral | 0.33 | Neutral | -1.443 |
| 119 | F403C | Disease | 0.87 | Disease | 0.906 | Disease | 0 | Disease | 0.725 | Disease | 0.828 | Disease | 0.72 | Deleterious | -6.957 |
| 120 | F403S | Disease | 0.748 | Disease | 0.892 | Disease | 0 | Disease | 0.69 | Disease | 0.746 | Disease | 0.72 | Deleterious | -7.007 |
| 121 | L406P | Disease | 0.799 | Disease | 0.934 | Disease | 0 | Disease | 0.755 | Disease | 0.877 | Disease | 0.66 | Deleterious | -6.675 |
| 122 | V407I | Neutral | 0.177 | Neutral | 0.359 | Neutral | 0.18 | Neutral | 0.29 | Disease | 0.428 | Neutral | 0.28 | Neutral | -0.888 |
| 123 | G413S | Disease | 0.553 | Disease | 0.745 | Disease | 0.01 | Disease | 0.675 | Disease | 0.616 | Neutral | 0.39 | Deleterious | -5.723 |
| 124 | E419G | Disease | 0.569 | Disease | 0.764 | Disease | 0.01 | Disease | 0.735 | Disease | 0.675 | Disease | 0.67 | Deleterious | -6.624 |
| 125 | E419K | Disease | 0.577 | Disease | 0.783 | Disease | 0.02 | Disease | 0.76 | Disease | 0.649 | Disease | 0.66 | Deleterious | -3.799 |
| 126 | E419D | Neutral | 0.221 | Disease | 0.513 | Neutral | 0.18 | Neutral | 0.375 | Disease | 0.463 | Neutral | 0.49 | Deleterious | -2.841 |
| 127 | D422H | Neutral | 0.439 | Disease | 0.696 | Disease | 0.03 | Disease | 0.675 | Disease | 0.737 | Neutral | 0.4 | Deleterious | -5.201 |
| 128 | D423N | Neutral | 0.295 | Neutral | 0.3 | Neutral | 0.43 | Neutral | 0.23 | Neutral | 0.422 | Neutral | 0.28 | Neutral | -2.124 |
| 129 | P426A | Neutral | 0.085 | Neutral | 0.127 | Neutral | 0.91 | Neutral | 0.265 | Neutral | 0.167 | Neutral | 0.33 | Deleterious | -4.291 |
| 130 | P426S | Neutral | 0.19 | Neutral | 0.243 | Neutral | 0.6 | Neutral | 0.255 | Neutral | 0.4 | Neutral | 0.36 | Deleterious | -4.371 |
| 131 | I428T | Neutral | 0.195 | Neutral | 0.417 | Neutral | 0.2 | Neutral | 0.305 | Neutral | 0.444 | Neutral | 0.34 | Deleterious | -2.857 |
| 132 | A429V | Neutral | 0.223 | Neutral | 0.365 | Neutral | 0.47 | Neutral | 0.385 | Neutral | 0.424 | Neutral | 0.3 | Neutral | -0.605 |
| 133 | M438T | Neutral | 0.239 | Disease | 0.562 | Disease | 0 | Disease | 0.66 | Disease | 0.737 | Disease | 0.65 | Deleterious | -4.063 |
| 134 | D439N | Disease | 0.574 | Neutral | 0.332 | Disease | 0 | Disease | 0.72 | Disease | 0.618 | Neutral | 0.37 | Deleterious | -4.848 |
| 135 | A440T | Neutral | 0.35 | Disease | 0.551 | Disease | 0.03 | Disease | 0.58 | Disease | 0.642 | Disease | 0.76 | Deleterious | -3.187 |
| 136 | S441L | Disease | 0.676 | Disease | 0.786 | Disease | 0.01 | Disease | 0.625 | Disease | 0.717 | Disease | 0.82 | Deleterious | -5.584 |
| 137 | I444V | Neutral | 0.215 | Neutral | 0.264 | Neutral | 0.1 | Neutral | 0.25 | Neutral | 0.456 | Neutral | 0.65 | Neutral | -0.691 |
| 138 | V447I | Neutral | 0.216 | Neutral | 0.162 | Neutral | 0.35 | Neutral | 0.215 | Neutral | 0.303 | Neutral | 0.33 | Neutral | -0.533 |
| 139 | R450C | NA | NA | Disease | 0.631 | Disease | 0 | Disease | 0.675 | Disease | 0.732 | Disease | 0.72 | Deleterious | -6.802 |
| 140 | R450H | NA | NA | Neutral | 0.41 | Disease | 0 | Disease | 0.615 | Disease | 0.334 | Disease | 0.65 | Deleterious | -3.778 |
| 141 | R450P | NA | NA | Disease | 0.806 | Disease | 0 | Disease | 0.775 | Disease | 0.772 | Disease | 0.79 | Deleterious | -5.848 |
| 142 | F451L | Neutral | 0.404 | Neutral | 0.499 | Disease | 0.01 | Disease | 0.67 | Disease | 0.508 | Disease | 0.76 | Deleterious | -5.24 |
| 143 | S453T | Neutral | 0.322 | Neutral | 0.394 | Neutral | 0.15 | Neutral | 0.3 | Neutral | 0.469 | Neutral | 0.4 | Neutral | -1.381 |
| 144 | I455V | Neutral | 0.292 | Neutral | 0.183 | Neutral | 0.52 | Neutral | 0.275 | Neutral | 0.254 | Neutral | 0.33 | Neutral | -0.078 |
| 145 | I456V | Neutral | 0.292 | Neutral | 0.389 | Neutral | 0.06 | Neutral | 0.325 | Neutral | 0.462 | Neutral | 0.56 | Neutral | -0.936 |
| 146 | L461V | Neutral | 0.388 | Disease | 0.685 | Neutral | 0.16 | Disease | 0.54 | Disease | 0.802 | Disease | 0.92 | Neutral | -2.434 |
| 147 | S462P | Disease | 0.639 | Disease | 0.884 | Disease | 0 | Disease | 0.605 | Disease | 0.752 | Disease | 0.88 | Deleterious | -4.903 |
| 148 | D465A | Disease | 0.544 | Disease | 0.537 | Disease | 0 | Disease | 0.675 | Disease | 0.553 | Neutral | 0.42 | Deleterious | -6.512 |
| 149 | D465T | Disease | 0.613 | Disease | 0.512 | Disease | 0 | Disease | 0.67 | Disease | 0.675 | Neutral | 0.44 | Deleterious | -5.748 |
| 150 | I466S | Disease | 0.634 | Neutral | 0.512 | Disease | 0 | Disease | 0.595 | Disease | 0.575 | Disease | 0.56 | Deleterious | -2.548 |
| 151 | P468L | Disease | 0.565 | Disease | 0.611 | Disease | 0 | Disease | 0.645 | Disease | 0.684 | Disease | 0.58 | Deleterious | -9.473 |
| 152 | H474T | Neutral | 0.11 | Neutral | 0.227 | Neutral | 0.63 | Neutral | 0.2 | Neutral | 0.399 | Neutral | 0.24 | Neutral | -2.477 |
| 153 | P475S | Neutral | 0.295 | Disease | 0.59 | Disease | 0.03 | Disease | 0.6 | Disease | 0.502 | Disease | 0.43 | Deleterious | -7.158 |
| 154 | V476I | Neutral | 0.423 | Neutral | 0.269 | Neutral | 0.1 | Neutral | 0.255 | Neutral | 0.468 | Neutral | 0.33 | Neutral | -0.853 |
| 155 | T477S | Neutral | 0.203 | Neutral | 0.154 | Neutral | 0.09 | Neutral | 0.15 | Neutral | 0.302 | Neutral | 0.4 | Neutral | -1.295 |
| 156 | T480A | Neutral | 0.146 | Neutral | 0.305 | Disease | 0.03 | Neutral | 0.36 | Neutral | 0.449 | Neutral | 0.33 | Neutral | -0.681 |
| 157 | M483I | Neutral | 0.198 | Neutral | 0.375 | Neutral | 0.06 | Neutral | 0.455 | Neutral | 0.459 | Disease | 0.7 | Deleterious | -3.456 |
| 158 | M483T | Disease | 0.66 | Disease | 0.593 | Disease | 0.01 | Disease | 0.74 | Disease | 0.759 | Disease | 0.9 | Deleterious | -5.617 |
| 159 | L485P | Disease | 0.881 | Disease | 0.839 | Disease | 0.01 | Disease | 0.695 | Disease | 0.772 | Disease | 0.77 | Deleterious | -6.865 |
| 160 | A486T | Neutral | 0.272 | Neutral | 0.325 | Neutral | 0.54 | Neutral | 0.15 | Neutral | 0.459 | Neutral | 0.45 | Neutral | -1.098 |
| 161 | R487W | Disease | 0.924 | Disease | 0.76 | Disease | 0 | Disease | 0.805 | Disease | 0.861 | Disease | 0.86 | Deleterious | -7.84 |
| 162 | V488A | Neutral | 0.105 | Neutral | 0.3 | Neutral | 0.46 | Neutral | 0.48 | Neutral | 0.437 | Neutral | 0.28 | Neutral | 0.758 |
| 163 | C489S | Neutral | 0.229 | Neutral | 0.413 | Neutral | 0.96 | Disease | 0.535 | Neutral | 0.465 | Neutral | 0.4 | Deleterious | -6.633 |
| 164 | M493L | Neutral | 0.124 | Neutral | 0.243 | Neutral | 1 | Neutral | 0.44 | Neutral | 0.428 | Neutral | 0.4 | Neutral | -0.656 |
| 165 | I494F | Disease | 0.669 | Disease | 0.804 | Disease | 0.04 | Disease | 0.585 | Disease | 0.635 | Disease | 0.57 | Deleterious | -3.92 |
| 166 | R497C | Disease | 0.838 | Disease | 0.793 | Disease | 0.01 | Disease | 0.755 | Disease | 0.834 | Disease | 0.48 | Deleterious | -6.613 |
| 167 | R497S | Neutral | 0.449 | Disease | 0.521 | Disease | 0.12 | Disease | 0.665 | Disease | 0.514 | Disease | 0.46 | Deleterious | -4.444 |
| 168 | R497I | Neutral | 0.625 | Disease | 589 | Disease | 0.1 | Disease | 0.76 | Disease | 0.77 | Disease | 0.49 | Deleterious | -6.58 |
| 169 | Q501P | Disease | 0.702 | Disease | 0.868 | Neutral | 0.08 | Disease | 0.665 | Disease | 0.514 | Disease | 0.74 | Deleterious | -5.78 |
| 170 | A503G | Neutral | 0.381 | Neutral | 0.321 | Neutral | 0.32 | Neutral | 0.31 | Neutral | 0.46 | Disease | 0.55 | Deleterious | -2.818 |
| 171 | A503T | Neutral | 0.193 | Neutral | 0.175 | Neutral | 0.35 | Neutral | 0.32 | Neutral | 0.307 | Neutral | 0.27 | Neutral | -1.567 |
| 172 | I504V | Neutral | 0.2 | Neutral | 0.234 | Neutral | 0.59 | Neutral | 0.175 | Neutral | 0.437 | Neutral | 0.28 | Neutral | -0.325 |
| 173 | F508L | Neutral | 0.35 | Disease | 0.743 | Disease | 0.1 | Disease | 0.665 | Disease | 0.668 | Disease | 0.84 | Deleterious | -5.418 |
| 174 | T510A | Neutral | 0.173 | Neutral | 0.229 | Neutral | 0.52 | Neutral | 0.295 | Neutral | 0.426 | Neutral | 0.54 | Neutral | -0.97 |
| 175 | R511W | Disease | 0.926 | Disease | 0.948 | Disease | 0 | Disease | 0.82 | Disease | 0.897 | Disease | 0.93 | Deleterious | -7.707 |
| 176 | D513V | NA | NA | Disease | 0.797 | Disease | 0 | Disease | 0.645 | Disease | 0.734 | Disease | 0.61 | Deleterious | -8.437 |
| 177 | I514T | NA | NA | Neutral | 0.384 | Neutral | 0.75 | Neutral | 320 | Neutral | 0.146 | Neutral | 0.33 | Neutral | 0.062 |
| 178 | I517M | Neutral | 0.494 | Neutral | 0.381 | Neutral | 0.07 | Neutral | 0.23 | Neutral | 0.414 | Neutral | 0.41 | Neutral | -1.971 |
| 179 | R518W | Disease | 0.902 | Disease | 0.792 | Disease | 0 | Disease | 0.61 | Disease | 0.772 | Disease | 0.87 | Deleterious | -7.54 |
| 180 | L523F | Disease | 0.517 | Disease | 0.598 | Disease | 0.01 | Disease | 0.555 | Disease | 0.513 | Neutral | 0.47 | Deleterious | -3.32 |
| 181 | V531F | Neutral | 0.487 | Disease | 0.908 | Disease | 0 | Disease | 0.665 | Disease | 0.775 | Disease | 0.89 | Deleterious | -4.467 |
| 182 | V531I | Neutral | 0 | Neutral | 0.303 | Neutral | 0.28 | Neutral | 0.29 | Neutral | 0.445 | Neutral | 0.49 | Neutral | -0.813 |
| 183 | G534D | Disease | 0.557 | Disease | 0.946 | Disease | 0 | Disease | 0.77 | Disease | 0.89 | Disease | 0.94 | Deleterious | -6.743 |
| 184 | V536M | Disease | 0.656 | Disease | 0.759 | Disease | 0 | Disease | 0.575 | Disease | 0.68 | Disease | 0.84 | Deleterious | -2.819 |
| 185 | F538S | Disease | 0.834 | Disease | 0.952 | Disease | 0 | Disease | 0.7 | Disease | 0.85 | Disease | 0.94 | Deleterious | -7.684 |
| 186 | Y542C | Disease | 0.943 | Disease | 0.944 | Disease | 0 | Disease | 0.83 | Disease | 0.897 | Disease | 0.94 | Deleterious | -8.57 |
| 187 | Q543P | Neutral | 0.281 | Disease | 0.718 | Neutral | 0.32 | Neutral | 0.33 | Disease | 0.559 | Neutral | 0.31 | Neutral | -1.45 |
| 188 | M545L | Neutral | 0.318 | Disease | 0.864 | Disease | 0.04 | Disease | 0.56 | Disease | 0.747 | Disease | 0.66 | Neutral | -1.488 |
| 189 | E546K | Disease | 0.503 | Disease | 0.746 | Neutral | 0.1 | Neutral | 0.465 | Disease | 0.567 | Neutral | 0.45 | Deleterious | -3.476 |
| 190 | E546Q | Disease | 0.52 | Disease | 0.578 | Neutral | 0.13 | Neutral | 210 | Neutral | 0.485 | Neutral | 0.46 | Deleterious | -2.69 |
| 191 | S547G | Neutral | 0.252 | Neutral | 0.486 | Neutral | 0.4 | Neutral | 0.275 | Neutral | 0.448 | Neutral | 0.38 | Neutral | -2.33 |
| 192 | A550V | Disease | 0.607 | Neutral | 0.413 | Disease | 0.05 | Neutral | 0.405 | Disease | 0.592 | Neutral | 0.47 | Deleterious | -3.239 |
| 193 | N564T | Neutral | 0.203 | Neutral | 0.181 | Neutral | 0.1 | Neutral | 0.395 | Neutral | 0.322 | Neutral | 0.49 | Deleterious | -2.755 |
| 194 | I569F | Disease | 0.634 | Disease | 0.511 | Neutral | 0.22 | Neutral | 0.255 | Neutral | 0.492 | Neutral | 0.61 | Deleterious | -3.82 |
| 195 | D573G | Neutral | 0.275 | Neutral | 0.482 | Disease | 0.1 | Disease | 0.61 | Neutral | 0.5 | Neutral | 0.66 | Deleterious | -6.539 |
| 196 | V579I | Neutral | 0.139 | Neutral | 0.035 | Neutral | 0.41 | Neutral | 0.08 | Neutral | 0.237 | Neutral | 0.23 | Neutral | 0.532 |
| 197 | A580T | Disease | 0.545 | Neutral | 0.229 | Disease | 0.03 | Disease | 0.54 | Disease | 0.503 | Disease | 0.76 | Deleterious | -3.853 |
| 198 | L581P | Disease | 0.881 | Disease | 0.863 | Neutral | 0.09 | Disease | 0.635 | Disease | 0.781 | Disease | 0.87 | Deleterious | -6.677 |
| 199 | E586V | Neutral | 0.359 | Neutral | 0.371 | Disease | 0.04 | Disease | 0.51 | Neutral | 0.489 | Neutral | 0.61 | Deleterious | -3.668 |
| 200 | C588F | Disease | 0.881 | Disease | 0.881 | Disease | 0.03 | Disease | 0.615 | Disease | 0.808 | Disease | 0.9 | Deleterious | -9.698 |
| 201 | E589K | Neutral | 0.256 | Neutral | 0.31 | Neutral | 0.08 | Neutral | 0.3 | Neutral | 0.458 | Neutral | 0.39 | Deleterious | -2.902 |
| 202 | R592C | Disease | 0.631 | Disease | 0.832 | Disease | 0 | Disease | 0.67 | Disease | 0.765 | Disease | 0.92 | Deleterious | -7.518 |
| 203 | R592H | Disease | 0.715 | Disease | 0.64 | Disease | 0.03 | Disease | 0.665 | Disease | 0.683 | Disease | 0.76 | Deleterious | -4.694 |
| 204 | R592P | Disease | 0.715 | Disease | 0.925 | Disease | 0.01 | Disease | 0.735 | Disease | 0.817 | Disease | 0.76 | Deleterious | -6.532 |
| 205 | G593R | Disease | 0.568 | Disease | 0.924 | Disease | 0.01 | Disease | 0.67 | Disease | 0.765 | Disease | 0.88 | Deleterious | -7.618 |
| 206 | I595M | Disease | 0.55 | Disease | 0.517 | Neutral | 0.08 | Neutral | 0.2 | Neutral | 0.49 | Neutral | 0.39 | Neutral | -1.964 |
| 207 | G602D | Neutral | 0.299 | Disease | 0.962 | Disease | 0 | Disease | 0.855 | Disease | 0.914 | Disease | 0.94 | Deleterious | -6.661 |
| 208 | G607A | Disease | 0.788 | Disease | 0.912 | Disease | 0 | Disease | 0.64 | Disease | 0.751 | Disease | 0.88 | Deleterious | -5.713 |
| 209 | I608M | Disease | 0.65 | Disease | 0.762 | Disease | 0.02 | Neutral | 0.405 | Disease | 0.606 | Disease | 0.79 | Neutral | -2.354 |
| 210 | V611A | Neutral | 0.137 | Neutral | 0.223 | Neutral | 0.08 | Neutral | 0.25 | Neutral | 0.39 | Neutral | 0.23 | Neutral | 0.929 |
| 211 | R616C | Disease | 0.922 | Disease | 0.967 | Disease | 0 | Disease | 0.81 | Disease | 0.909 | Disease | 0.94 | Deleterious | -7.518 |
| 212 | R616P | Disease | 0.806 | Disease | 0.973 | Disease | 0 | Disease | 0.8 | Disease | 0.915 | Disease | 0.94 | Deleterious | -6.532 |
| 213 | R616Q | Disease | 0.531 | Disease | 0.946 | Disease | 0 | Disease | 0.755 | Disease | 0.885 | Disease | 0.94 | Deleterious | -3.709 |
| 214 | R616W | Disease | 0.93 | Disease | 0.973 | Disease | 0 | Disease | 0.805 | Disease | 0.894 | Disease | 0.89 | Deleterious | -7.518 |
| 215 | V618I | Neutral | 0.404 | Disease | 0.797 | Disease | 0.02 | Neutral | 0.49 | Disease | 0.57 | Disease | 0.76 | Neutral | -0.952 |
| 216 | I619V | Neutral | 0.212 | Neutral | 0.446 | Neutral | 0.52 | Neutral | 0.165 | Neutral | 0.464 | Neutral | 0.33 | Neutral | -0.751 |
| 217 | G622V | Disease | 0.903 | Disease | 0.976 | Disease | 0 | Disease | 0.72 | Disease | 0.841 | Disease | 0.94 | Deleterious | -8.526 |
| 218 | V623I | Neutral | 0.102 | Neutral | 0.136 | Neutral | 0.71 | Neutral | 0.15 | Neutral | 0.27 | Neutral | 0.31 | Neutral | 0.288 |
| 219 | V626I | Neutral | 0.175 | Neutral | 0.396 | Neutral | 0.43 | Neutral | 0.22 | Neutral | 0.454 | Neutral | 0.47 | Neutral | -0.819 |
| 220 | T628I | Neutral | 0.422 | Neutral | 0.482 | Disease | 0.03 | Disease | 0.56 | Neutral | 0.492 | Disease | 0.71 | Deleterious | -5.71 |
| 221 | Q629H | Neutral | 0.371 | Neutral | 0.409 | Neutral | 0.12 | Neutral | 0.235 | Neutral | 0.462 | Neutral | 0.25 | Deleterious | -3.281 |
| 222 | R631H | Disease | 0.706 | Disease | 0.612 | Neutral | 0.06 | Neutral | 0.29 | Disease | 0.521 | Disease | 0.57 | Deleterious | -4.557 |
| 223 | R631L | Disease | 0.7 | Disease | 0.805 | Neutral | 0.2 | Neutral | 0.49 | Disease | 0.639 | Disease | 0.59 | Deleterious | -6.383 |
| 224 | R631P | Disease | 0.801 | Disease | 0.654 | Neutral | 0.21 | Disease | 0.565 | Disease | 0.589 | Disease | 0.51 | Deleterious | -6.284 |
| 225 | I632C | Disease | 0.827 | Disease | 0.742 | Disease | 0.02 | Disease | 0.52 | Disease | 0.645 | Disease | 0.92 | Deleterious | -4.399 |
| 226 | K634E | Neutral | 0.423 | Disease | 0.585 | Neutral | 0.08 | Neutral | 0.305 | Neutral | 0.49 | Neutral | 0.62 | Deleterious | -2.927 |
| 227 | A635V | Disease | 0.622 | Disease | 0.624 | Disease | 0.01 | Disease | 0.545 | Disease | 0.653 | Disease | 0.72 | Deleterious | -3.762 |
| 228 | R647L | Disease | 0.522 | Disease | 0.699 | Neutral | 0.24 | Disease | 0.525 | Disease | 0.613 | Disease | 0.61 | Deleterious | -5.51 |
| 229 | F654L | Neutral | 0.311 | Neutral | 0.474 | Neutral | 0.12 | Disease | 0.585 | Disease | 0.836 | Neutral | 0.4 | Deleterious | -5.463 |
| 230 | R658H | Disease | 0.87 | Disease | 0.88 | Disease | 0.02 | Disease | 0.69 | Disease | 0.808 | Disease | 0.61 | Deleterious | -4.563 |
| 231 | R658G | Disease | 0.831 | Disease | 0.908 | Disease | 0.02 | Disease | 0.705 | Disease | 0.791 | Disease | 0.85 | Deleterious | -6.38 |
| 232 | H659Y | Disease | 0.589 | Disease | 0.785 | Neutral | 0.34 | Neutral | 0.245 | Disease | 0.683 | Disease | 0.84 | Deleterious | -4.304 |
| 233 | A660S | Neutral | 0.454 | Disease | 0.721 | Neutral | 0.06 | Neutral | 0.415 | Disease | 0.548 | Disease | 0.57 | Neutral | -1.938 |
| 234 | C663R | Disease | 0.793 | Disease | 0.934 | Disease | 0.02 | Disease | 0.765 | Disease | 0.871 | Disease | 0.91 | Deleterious | -11.243 |
| 235 | R666Q | Disease | 0.772 | Disease | 0.935 | Disease | 0 | Disease | 0.84 | Disease | 0.903 | Disease | 0.94 | Deleterious | -3.737 |
| 236 | R666W | Disease | 0.959 | Disease | 0.956 | Disease | 0 | Disease | 0.84 | Disease | 0.901 | Disease | 0.94 | Deleterious | -7.474 |
| 237 | T672A | Neutral | 0.158 | Neutral | 0.473 | Neutral | 0.63 | Neutral | 0.37 | Neutral | 0.431 | Neutral | 0.54 | Neutral | -2.488 |
| 238 | T672M | Neutral | 0.482 | Disease | 0.506 | Neutral | 0.06 | Neutral | 0.26 | Neutral | 0.496 | Neutral | 0.63 | Deleterious | -3.787 |
| 239 | G675S | Disease | 0.808 | Disease | 0.805 | Disease | 0.01 | Disease | 0.615 | Disease | 0.664 | Disease | 0.76 | Deleterious | -5.609 |
| 240 | M677I | Disease | 0.503 | Neutral | 0.254 | Neutral | 0.11 | Neutral | 0.315 | Neutral | 0.225 | Neutral | 0.46 | Deleterious | -3.74 |
| 241 | M677V | Neutral | 0.481 | Disease | 0.587 | Neutral | 0.18 | Neutral | 0.35 | Neutral | 0.493 | Neutral | 0.48 | Deleterious | -3.74 |
| 242 | M677T | Disease | 0.55 | Disease | 0.75 | Disease | 0.02 | Disease | 0.515 | Disease | 0.656 | Disease | 0.77 | Deleterious | -5.609 |
| 243 | V678A | Disease | 0.523 | Disease | 0.592 | Disease | 0.04 | Disease | 0.533 | Disease | 0.641 | Disease | 0.92 | Deleterious | -3.488 |
| 244 | D681N | Disease | 0.738 | Disease | 0.94 | Disease | 0 | Disease | 0.83 | Disease | 0.928 | Disease | 0.93 | Deleterious | -4.674 |
| 245 | K682T | Neutral | 0.358 | Neutral | 0.361 | Disease | 0.03 | Disease | 0.6 | Neutral | 0.499 | Neutral | 0.7 | Deleterious | -5.464 |
| 246 | R683Q | Disease | 0.772 | Disease | 0.919 | Disease | 0 | Disease | 0.865 | Disease | 0.866 | Disease | 0.94 | Deleterious | -3.662 |
| 247 | R683W | Disease | 0.959 | Disease | 0.945 | Disease | 0 | Disease | 0.865 | Disease | 0.897 | Disease | 0.94 | Deleterious | -7.312 |
| 248 | F684C | Disease | 0.733 | Disease | 0.921 | Disease | 0 | Disease | 0.67 | Disease | 0.772 | Disease | 0.83 | Deleterious | -6.418 |
| 249 | F684L | Neutral | 0.325 | Disease | 0.831 | Disease | 0.01 | Neutral | 0.485 | Disease | 0.599 | Disease | 0.61 | Deleterious | -4.865 |
| 250 | R686H | Disease | 0.578 | Neutral | 0.404 | Disease | 0.01 | Disease | 0.535 | Disease | 0.623 | Neutral | 0.38 | Deleterious | -4.348 |
| 251 | R686C | Disease | 0.812 | Disease | 0.673 | Disease | 0 | Disease | 0.63 | Disease | 0.73 | Disease | 0.5 | Deleterious | -6.97 |
| 252 | R686P | Disease | 0.651 | Disease | 0.834 | Disease | 0.01 | Disease | 0.615 | Disease | 0.775 | Disease | 0.78 | Deleterious | -6.041 |
| 253 | R690P | Disease | 0.719 | Disease | 0.855 | Disease | 0 | Disease | 0.795 | Disease | 0.821 | Disease | 0.71 | Deleterious | -5.529 |
| 254 | R690Q | Neutral | 0.435 | Neutral | 0.31 | Disease | 0.03 | Disease | 0.605 | Neutral | 0.499 | Neutral | 0.39 | Deleterious | -3.042 |
| 255 | R690W | Disease | 0.891 | Disease | 0.741 | Disease | 0 | Disease | 0.71 | Disease | 0.785 | Disease | 0.52 | Deleterious | -6.665 |
| 256 | K692N | Disease | 0.597 | Disease | 0.578 | Disease | 0 | Disease | 0.685 | Disease | 0.538 | Neutral | 0.43 | Deleterious | -4.539 |
| 257 | P694S | Disease | 0.748 | Neutral | 0.284 | Neutral | 0.11 | Neutral | 0.415 | Disease | 0.509 | Neutral | 0.43 | Deleterious | -7.379 |
| 258 | R695H | Neutral | 0.303 | Neutral | 0.265 | Neutral | 0.09 | Neutral | 0.4 | Neutral | 0.447 | Neutral | 0.38 | Deleterious | -2.766 |
| 259 | R695P | Neutral | 0.134 | Disease | 0.723 | Neutral | 0.27 | Neutral | 0.45 | Disease | 0.574 | Neutral | 0.49 | Deleterious | -3.371 |
| 260 | R695S | Neutral | 0.168 | Neutral | 0.237 | Neutral | 0.08 | Neutral | 0.465 | Neutral | 0.437 | Neutral | 0.32 | Neutral | -2.258 |
| 261 | H700Y | Neutral | 0.159 | Neutral | 0.306 | Neutral | 1 | Neutral | 0.26 | Neutral | 0.454 | Neutral | 0.34 | Neutral | 0.776 |
| 262 | T702I | Neutral | 0.278 | Neutral | 0.306 | Neutral | 0.21 | Neutral | 0.33 | Neutral | 0.415 | Neutral | 0.4 | Deleterious | -3.175 |
| 263 | A704V | Neutral | 0.32 | Neutral | 0.297 | Disease | 0.04 | Disease | 0.55 | Neutral | 0.429 | Neutral | 0.31 | Neutral | -2.299 |
| 264 | N705D | Neutral | 0.182 | Neutral | 0.274 | Neutral | 0.69 | Neutral | 0.23 | Neutral | 0.404 | Neutral | 0.28 | Neutral | -1.324 |
| 265 | N705I | Neutral | 0.258 | Disease | 0.588 | Neutral | 0.4 | Neutral | 0.395 | Disease | 0.534 | Disease | 0.56 | Neutral | -0.209 |
| 266 | N705S | Neutral | 0.171 | Neutral | 0.287 | Neutral | 0.87 | Neutral | 0.23 | Neutral | 0.404 | Neutral | 0.25 | Neutral | -0.197 |
| 267 | N707T | Disease | 0.687 | Disease | 0.681 | Neutral | 0.19 | Neutral | 0.395 | Disease | 0.561 | Disease | 0.56 | Deleterious | -5.097 |
| 268 | T709I | Disease | 0.508 | Disease | 0.515 | Neutral | 0.11 | Neutral | 0.445 | Neutral | 0.488 | Neutral | 0.44 | Deleterious | -3.048 |
| 269 | V710M | Neutral | 0.48 | Disease | 0.824 | Disease | 0.02 | Disease | 0.53 | Disease | 0.608 | Neutral | 0.49 | Neutral | -1.366 |
| 270 | D711N | Neutral | 0.345 | Disease | 0.789 | Disease | 0.04 | Neutral | 0.455 | Disease | 0.578 | Neutral | 0.43 | Deleterious | -3.574 |
| 271 | D711Y | Disease | 0.748 | Disease | 0.922 | Disease | 0 | Disease | 0.67 | Disease | 0.77 | Disease | 0.61 | Deleterious | -6.481 |
| 272 | G713R | Disease | 0.504 | Disease | 0.793 | Disease | 0.01 | Disease | 0.695 | Disease | 0.789 | Disease | 0.67 | Neutral | -1.204 |
| 273 | A717G | Neutral | 0.428 | Disease | 0.631 | Neutral | 0.08 | Disease | 0.54 | Disease | 0.336 | Disease | 0.71 | Neutral | -2.489 |
| 274 | R722P | Disease | 0.573 | Disease | 0.919 | Neutral | 0.2 | Disease | 0.72 | Disease | 0.777 | Disease | 0.79 | Deleterious | -5.422 |
| 275 | M724T | Disease | 0.684 | Disease | 0.823 | Disease | 0.01 | Disease | 0.765 | Disease | 0.781 | Disease | 0.81 | Deleterious | -5.151 |
| 276 | A725P | Disease | 0.623 | Disease | 0.867 | Disease | 0.04 | Disease | 0.73 | Disease | 0.669 | Disease | 0.55 | Deleterious | -4.097 |

### Assessment of the functional impact of mutations

The mutation assessor predicted 57 mutations as high impacting on protein functionality out of 276 mutations screened [Table 2].

**Table 2.** Prediction of Functional Impact using MutationAssessor Server

| Sr.no | Mutants | MutationAssessor Prediction |  |  |  |
| --- | --- | --- | --- | --- | --- |
|  |  | FI score | VC score | VS score | Functional Impact |
| 1 | E20K | 3.785 | 4.57 | 3 | high |
| 2 | Q21E | 3.855 | 4.71 | 3 | high |
| 3 | S23F | 2.865 | 3.43 | 2.3 | medium |
| 4 | M25L | 3.69 | 4.24 | 3.14 | high |
| 5 | R26Q | 1.63 | 3.26 | 0 | low |
| 6 | R26W | 0.185 | 1.06 | -0.69 | neutral |
| 7 | R30C | 3.525 | 4.16 | 2.89 | high |
| 8 | T31M | 3.265 | 4.23 | 2.3 | medium |
| 9 | H37R | 3.61 | 4.08 | 3.14 | high |
| 10 | V39I | 1.265 | 0.13 | 2.4 | low |
| 11 | G47R | 3.97 | 4.8 | 3.14 | high |
| 12 | K48R | 4.315 | 5.49 | 3.14 | high |
| 13 | T49I | 4.25 | 5.36 | 3.14 | high |
| 14 | L55P | 3.17 | 3.25 | 3.09 | medium |
| 15 | M57K | 1.04 | 1.39 | 0.69 | low |
| 16 | P64L | 3.19 | 3.24 | 3.14 | medium |
| 17 | E66V | 2.005 | 2.62 | 1.39 | medium |
| 18 | R75I | 4.46 | 5.62 | 3.3 | high |
| 19 | T76A | 4.46 | 5.62 | 3.3 | high |
| 20 | P78S | 1.225 | -0.81 | 3.26 | low |
| 21 | E86Q | 3.265 | 3.27 | 3.26 | medium |
| 22 | R88G | 2.73 | 2.9 | 2.56 | medium |
| 23 | N92S | 0.55 | 0 | 1.1 | neutral |
| 24 | F93L | 0.77 | 1.13 | 0.41 | neutral |
| 25 | Y94S | 2.61 | 2.22 | 3 | medium |
| 26 | E95D | 2.175 | 1.41 | 2.94 | medium |
| 27 | Q97L | 1.195 | 1.14 | 1.25 | low |
| 28 | E98D | -0.39 | 0.32 | -1.1 | neutral |
| 29 | K101T | 1.085 | 1.07 | 1.1 | low |
| 30 | P103L | 1.83 | 2.97 | 0.69 | low |
| 31 | P103R | 0.61 | 0.53 | 0.69 | neutral |
| 32 | R112C | 4.44 | 5.62 | 3.26 | high |
| 33 | R112G | 4.44 | 5.62 | 3.26 | high |
| 34 | R112H | 4.44 | 5.62 | 3.26 | high |
| 35 | H118R | 3.35 | 3.81 | 2.89 | medium |
| 36 | P123T | -0.185 | -0.37 | 0 | neutral |
| 37 | R125H | 2.72 | 2.26 | 3.18 | medium |
| 38 | D131N | 1.88 | 0.5 | 3.26 | low |
| 39 | C134T | NA | NA | NA | NA |
| 40 | H135D | 1.91 | 1.42 | 2.4 | low |
| 41 | S136N | 1.785 | 1.27 | 2.3 | low |
| 42 | Y141C | 2.31 | 2.14 | 2.48 | medium |
| 43 | V142L | 2.22 | 1.61 | 2.83 | medium |
| 44 | R143Q | 3.3 | 3.38 | 3.22 | medium |
| 45 | R143G | 3.3 | 3.38 | 3.22 | medium |
| 46 | R143T | NA | NA | NA | NA |
| 47 | A144V | 2.04 | 2 | 2.08 | medium |
| 48 | H148T | NA | NA | NA | NA |
| 49 | D149N | 1.135 | 0.93 | 1.34 | low |
| 51 | L152R | 1.735 | 2.08 | 1.39 | low |
| 52 | Y158C | 3.405 | 4.04 | 2.77 | medium |
| 53 | H164R | 0.76 | -0.09 | 1.61 | neutral |
| 54 | E167Q | 0.85 | 0.6 | 1.1 | low |
| 55 | V168M | 1.655 | 1.7 | 1.61 | low |
| 56 | L170F | 2.365 | 1.79 | 2.94 | medium |
| 57 | L170R | 3.145 | 3.35 | 2.94 | medium |
| 58 | A172T | 0.94 | 1.37 | 0.51 | low |
| 59 | N176S | 0.17 | -0.41 | 0.75 | neutral |
| 60 | D178E | 0.255 | -1.48 | 1.99 | neutral |
| 60 | G184V | 4.365 | 5.47 | 3.26 | high |
| 61 | R185T | NA | NA | NA | NA |
| 62 | R186C | 2.675 | 3.74 | 1.61 | medium |
| 63 | R186H | 2.125 | 2.64 | 1.61 | medium |
| 64 | Q187R | -0.985 | -0.77 | -1.2 | neutral |
| 65 | S198L | -0.375 | -1.44 | 0.69 | neutral |
| 66 | I199M | 1.605 | 0.91 | 2.3 | low |
| 67 | H201T | NA | NA | NA | NA |
| 68 | A202V | 2.92 | 2.54 | 3.3 | medium |
| 69 | N203S | 2.22 | 2.3 | 2.14 | medium |
| 70 | Y209C | 4.47 | 5.61 | 3.33 | high |
| 71 | P215S | 3.115 | 2.9 | 3.33 | medium |
| 72 | A218V | 3.86 | 4.39 | 3.33 | high |
| 73 | L220M | 2.135 | 1.44 | 2.83 | medium |
| 74 | L220V | 0.665 | 0.69 | 0.64 | neutral |
| 75 | S222A | 3.27 | 3.24 | 3.3 | medium |
| 76 | A226T | 0.64 | 0.94 | 0.34 | neutral |
| 77 | R227C | 2.98 | 3.4 | 2.56 | medium |
| 78 | R227H | 2.98 | 3.4 | 2.56 | medium |
| 79 | K228T | 1.065 | 1.03 | 1.1 | low |
| 80 | A229V | 2.595 | 3.24 | 1.95 | medium |
| 81 | V231M | 4.315 | 5.3 | 3.33 | high |
| 82 | N241I | 3.92 | 4.58 | 3.26 | high |
| 83 | I244M | 3.645 | 4.03 | 3.26 | high |
| 84 | I244T | 3.445 | 3.63 | 3.26 | medium |
| 85 | D245G | 3.455 | 4.2 | 2.71 | medium |
| 86 | M247T | 2.43 | 1.72 | 3.14 | medium |
| 87 | N250S | 0.295 | 0.08 | 0.51 | neutral |
| 88 | T252P | 2.42 | 2.36 | 2.48 | medium |
| 89 | R253C | 3.37 | 3.91 | 2.83 | medium |
| 90 | R254G | 3.005 | 3.07 | 2.94 | medium |
| 91 | R258T | NA | NA | NA | NA |
| 92 | C259T | NA | NA | NA | NA |
| 93 | K274E | 2.485 | 3.36 | 1.61 | medium |
| 94 | Q279P | 2.02 | 2.94 | 1.1 | medium |
| 95 | R282Q | -0.56 | -1.12 | 0 | neutral |
| 96 | D283V | 1.775 | 2.86 | 0.69 | low |
| 97 | R286T | NA | NA | NA | NA |
| 98 | R287Y | NA | NA | NA | NA |
| 99 | A297V | -0.32 | -0.64 | 0 | neutral |
| 100 | R299T | NA | NA | NA | NA |
| 101 | E300Q | 1.245 | 1.1 | 1.39 | low |
| 102 | T301K | 2.125 | 2.64 | 1.61 | medium |
| 103 | T301M | 2.47 | 3.33 | 1.61 | medium |
| 104 | N307K | 2.68 | 3.75 | 1.61 | medium |
| 105 | D312Y | 2.8 | 3.99 | 1.61 | medium |
| 106 | V314A | 2.01 | 3.33 | 0.69 | medium |
| 107 | Q316E | 1.355 | 1.1 | 1.61 | low |
| 108 | Q316K | 0.645 | -0.32 | 1.61 | neutral |
| 109 | I323M | 2.8 | 3.99 | 1.61 | medium |
| 110 | R324H | 3.175 | 4.74 | 1.61 | medium |
| 111 | T325M | 1.545 | 2.4 | 0.69 | low |
| 112 | E327G | 2.985 | 4.58 | 1.39 | medium |
| 113 | R335W | 3.115 | 4.62 | 1.61 | medium |
| 114 | R345H | 2.8 | 3.99 | 1.61 | medium |
| 115 | F394L | 1.34 | 2.27 | 0.41 | low |
| 116 | P396L | 2.485 | 3.58 | 1.39 | medium |
| 117 | P396S | 0.965 | 0.54 | 1.39 | low |
| 118 | T398I | 1.01 | 0.92 | 1.1 | low |
| 119 | F403C | 3.125 | 4.64 | 1.61 | medium |
| 120 | F403S | 3.125 | 4.64 | 1.61 | medium |
| 121 | L406P | 3.16 | 4.71 | 1.61 | medium |
| 122 | V407I | 1.74 | 1.87 | 1.61 | low |
| 123 | G413S | 3.175 | 4.74 | 1.61 | medium |
| 124 | E419G | 3.64 | 4.51 | 2.77 | high |
| 125 | E419K | 3.64 | 4.51 | 2.77 | high |
| 126 | E419D | 2.49 | 2.21 | 2.77 | medium |
| 127 | D422H | 3.16 | 3.76 | 2.56 | medium |
| 128 | D423N | 1.81 | 1.61 | 2.01 | low |
| 129 | P426A | 1.555 | 0.34 | 2.77 | low |
| 130 | P426S | 2.155 | 1.54 | 2.77 | medium |
| 131 | I428T | 2.27 | 2.34 | 2.2 | medium |
| 132 | A429V | 1.1 | 1.1 | 1.1 | low |
| 133 | M438T | 2.67 | 3.04 | 2.3 | medium |
| 134 | D439N | 3.34 | 3.91 | 2.77 | medium |
| 135 | A440T | 3.22 | 4.36 | 2.08 | medium |
| 136 | S441L | 3.24 | 3.77 | 2.71 | medium |
| 137 | I444V | 2.25 | 2.8 | 1.7 | medium |
| 138 | V447I | 1.445 | 1.35 | 1.54 | low |
| 139 | R450C | 3.52 | 4.33 | 2.71 | high |
| 140 | R450H | 2.825 | 3.64 | 2.01 | medium |
| 141 | R450P | 3.52 | 4.33 | 2.71 | high |
| 142 | F451L | 3.58 | 4.45 | 2.71 | high |
| 143 | S453T | 1.33 | 1.78 | 0.88 | low |
| 144 | I455V | 0.015 | 0.39 | -0.36 | neutral |
| 145 | I456V | 2.825 | 2.94 | 2.71 | medium |
| 146 | L461V | 3.435 | 4.16 | 2.71 | medium |
| 147 | S462P | 3.675 | 4.58 | 2.77 | high |
| 148 | D465A | 3.465 | 4.16 | 2.77 | medium |
| 149 | D465T | NA | NA | NA | NA |
| 150 | I466S | 2.05 | 2.71 | 1.39 | medium |
| 151 | P468L | 3.665 | 4.56 | 2.77 | high |
| 152 | H474T | NA | NA | NA | NA |
| 153 | P475S | 3.15 | 3.53 | 2.77 | medium |
| 154 | V476I | 2.74 | 2.77 | 2.71 | medium |
| 155 | T477S | 0.6 | 0.69 | 0.51 | neutral |
| 156 | T480A | 0.785 | 0.18 | 1.39 | neutral |
| 157 | M483I | 1.99 | 2.44 | 1.54 | medium |
| 158 | M483T | 3.58 | 4.52 | 2.64 | high |
| 159 | L485P | 3.63 | 4.49 | 2.77 | high |
| 160 | A486T | 1.69 | 0.67 | 2.71 | low |
| 161 | R487W | 3.685 | 4.6 | 2.77 | high |
| 162 | V488A | 1.1 | 1.1 | 1.1 | low |
| 163 | C489S | 0.975 | 0.77 | 1.18 | low |
| 164 | M493L | 0.78 | 0.17 | 1.39 | neutral |
| 165 | I494F | 3.055 | 3.47 | 2.64 | medium |
| 166 | R497C | 2.9 | 3.5 | 2.3 | medium |
| 167 | R497S | 3.245 | 4.19 | 2.3 | medium |
| 168 | R497I | NA | NA | NA | NA |
| 169 | Q501P | 3.605 | 4.5 | 2.71 | high |
| 170 | A503G | 2.66 | 3.02 | 2.3 | medium |
| 171 | A503T | 1.23 | 1.77 | 0.69 | low |
| 172 | I504V | 1.48 | 0.76 | 2.2 | low |
| 173 | F508L | 3.48 | 4.48 | 2.48 | medium |
| 174 | T510A | 1.59 | 0.88 | 2.3 | low |
| 175 | R511W | 3.66 | 4.61 | 2.71 | high |
| 176 | D513V | 3.63 | 4.49 | 2.77 | high |
| 177 | I514T | -0.4 | -0.29 | -0.51 | neutral |
| 178 | I517M | 1.605 | 2.01 | 1.2 | low |
| 179 | R518W | 3.59 | 4.47 | 2.71 | high |
| 180 | L523F | 3.165 | 3.69 | 2.64 | medium |
| 181 | V531F | 4.2 | 5.36 | 3.04 | high |
| 182 | V531I | 1.44 | 0.93 | 1.95 | low |
| 183 | G534D | 4.615 | 5.83 | 3.4 | high |
| 184 | V536M | 4.3 | 5.27 | 3.33 | high |
| 185 | F538S | 4.61 | 5.82 | 3.4 | high |
| 186 | Y542C | 4.6 | 5.77 | 3.43 | high |
| 187 | Q543P | -0.265 | -0.53 | 0 | neutral |
| 188 | M545L | 0.475 | -0.11 | 1.06 | neutral |
| 189 | E546K | 2.2 | 1.73 | 2.67 | medium |
| 190 | E546Q | 2.5 | 1.63 | 3.37 | medium |
| 191 | S547G | 2.215 | 2.08 | 2.35 | medium |
| 192 | A550V | 2.69 | 3.08 | 2.3 | medium |
| 193 | N564T | 1.86 | 1.32 | 2.4 | low |
| 194 | I569F | 1.9 | 1.24 | 2.56 | low |
| 195 | D573G | 2.03 | 0.66 | 3.4 | medium |
| 196 | V579I | -0.17 | 0 | -0.34 | neutral |
| 197 | A580T | 2.325 | 1.22 | 3.43 | medium |
| 198 | L581P | 4.43 | 5.46 | 3.4 | high |
| 199 | E586V | 2.39 | 2.83 | 1.95 | medium |
| 200 | C588F | 3.955 | 4.48 | 3.43 | high |
| 201 | E589K | 1.095 | 0.24 | 1.95 | low |
| 202 | R592C | 3.315 | 3.26 | 3.37 | medium |
| 203 | R592H | 3.2 | 3.03 | 3.37 | medium |
| 204 | R592P | 3.315 | 3.26 | 3.37 | medium |
| 205 | G593R | 4.645 | 5.82 | 3.47 | high |
| 206 | I595M | 2.21 | 2.02 | 2.4 | medium |
| 207 | G602D | 4.265 | 5.1 | 3.43 | high |
| 208 | G607A | 4.29 | 5.15 | 3.43 | high |
| 209 | I608M | 3.13 | 3.86 | 2.4 | medium |
| 210 | V611A | -1.115 | -3.33 | 1.1 | neutral |
| 211 | R616C | 4.635 | 5.84 | 3.43 | high |
| 212 | R616P | 4.635 | 5.84 | 3.43 | high |
| 213 | R616Q | 4.285 | 5.14 | 3.43 | high |
| 214 | R616W | 4.635 | 5.84 | 3.43 | high |
| 215 | V618I | 3.04 | 2.65 | 3.43 | medium |
| 216 | I619V | 0.825 | 0.04 | 1.61 | low |
| 217 | G622V | 4.65 | 5.83 | 3.47 | high |
| 218 | V623I | -0.58 | -0.83 | -0.33 | neutral |
| 219 | V626I | 1.44 | 1.56 | 1.32 | low |
| 220 | T628I | 1.755 | 0.04 | 3.47 | low |
| 221 | Q629H | 1.425 | 0.37 | 2.48 | low |
| 222 | R631H | 2.445 | 1.52 | 3.37 | medium |
| 223 | R631L | 1.815 | 0.96 | 2.67 | low |
| 224 | R631P | 1.415 | -0.54 | 3.37 | low |
| 225 | I632C | NA | NA | NA | NA |
| 226 | K634E | 3.12 | 3.15 | 3.09 | medium |
| 227 | A635V | 2.25 | 1.79 | 2.71 | medium |
| 228 | R647L | 2.14 | 1.34 | 2.94 | medium |
| 229 | F654L | 2.22 | 1.01 | 3.43 | medium |
| 230 | R658H | 3.565 | 3.76 | 3.37 | high |
| 231 | R658G | 4.46 | 5.55 | 3.37 | high |
| 232 | H659Y | 3.22 | 3.3 | 3.14 | medium |
| 233 | A660S | 2.43 | 2.42 | 2.44 | medium |
| 234 | C663R | 4.02 | 4.64 | 3.4 | high |
| 235 | R666Q | 4.6 | 5.83 | 3.37 | high |
| 236 | R666W | 4.6 | 5.83 | 3.37 | high |
| 237 | T672A | 1.2 | 0.45 | 1.95 | low |
| 238 | T672M | 2.175 | 2.4 | 1.95 | medium |
| 239 | G675S | 2.43 | 2.15 | 2.71 | medium |
| 240 | M677I | 1.14 | -1.09 | 3.37 | low |
| 241 | M677V | 1.24 | -0.19 | 2.67 | low |
| 242 | M677T | 3.75 | 4.13 | 3.37 | high |
| 243 | V678A | 3.27 | 3.71 | 2.83 | medium |
| 244 | D681N | 4.255 | 5.11 | 3.4 | high |
| 245 | K682T | 2.54 | 1.99 | 3.09 | medium |
| 246 | R683Q | 4.59 | 5.81 | 3.37 | high |
| 247 | R683W | 4.59 | 5.81 | 3.37 | high |
| 248 | F684C | 4.095 | 5.05 | 3.14 | high |
| 249 | F684L | 2.94 | 3.44 | 2.44 | medium |
| 250 | R686H | 2.5 | 1.96 | 3.04 | medium |
| 251 | R686C | 3.74 | 4.44 | 3.04 | high |
| 252 | R686P | 3.19 | 3.34 | 3.04 | medium |
| 253 | R690P | 3.555 | 4.07 | 3.04 | high |
| 254 | R690Q | 1.395 | 0.44 | 2.35 | low |
| 255 | R690W | 3.005 | 2.97 | 3.04 | medium |
| 256 | K692N | 2.465 | 2.53 | 2.4 | medium |
| 257 | P694S | 2.06 | 0.69 | 3.43 | medium |
| 258 | R695H | 1.685 | 1.29 | 2.08 | low |
| 259 | R695P | 1.57 | 1.06 | 2.08 | low |
| 260 | R695S | 0.89 | -0.3 | 2.08 | low |
| 261 | H700Y | -0.245 | 0.47 | -0.96 | neutral |
| 262 | T702I | 1.62 | 2.14 | 1.1 | low |
| 263 | A704V | 1.53 | 2.37 | 0.69 | low |
| 264 | N705D | 0.25 | -0.6 | 1.1 | neutral |
| 265 | N705I | 1.965 | 2.83 | 1.1 | medium |
| 266 | N705S | 0.075 | 0.44 | -0.29 | neutral |
| 267 | N707T | 2.9 | 4.01 | 1.79 | medium |
| 268 | T709I | 2.085 | 2.56 | 1.61 | medium |
| 269 | V710M | 1.665 | 2.64 | 0.69 | low |
| 270 | D711N | 3.11 | 4.61 | 1.61 | medium |
| 271 | D711Y | 3.11 | 4.61 | 1.61 | medium |
| 272 | G713R | 0 | 0 | 0 | neutral |
| 273 | A717G | 2.54 | 4.39 | 0.69 | medium |
| 274 | R722P | 2.61 | 3.43 | 1.79 | medium |
| 275 | M724T | 2.545 | 3.99 | 1.1 | medium |
| 276 | A725P | 3.215 | 4.64 | 1.79 | medium |

### Protein stability analysis

mCSM, SDM, DUET, and STRUM were used to predict the impact of the mutation on protein stability. Different algorithms calculated varied results as illustrated in [Table 3]. mCSM predicted 245 mutations as destabilizing whereas SDM, DUET, and STRUM predicted 183, 212, and 68 mutations respectively as having the destabilizing effect on the protein.

**Table 3.** Protein Stability prediction using computational servers

| Protein Stability Prediction |  |  |  |  |  |  |  |  |
| --- | --- | --- | --- | --- | --- | --- | --- | --- |
| Sr.no | mCSM |  | SDM |  | DUET |  | STRUM |  |
| | $\Delta\Delta G$ | Prediction | $\Delta\Delta G$ | Prediction | $\Delta\Delta G$ | Prediction | $\Delta\Delta G$ | Prediction |
| 1 | -0.782 | Destabilizing | -1.43 | Destabilizing | -0.732 | Destabilizing | 0.96 | Stabilizing |
| 2 | -1.331 | Destabilizing | 0.35 | Stabilizing | -0.875 | Destabilizing | 2.17 | Stabilizing |
| 3 | -0.87 | Destabilizing | 0.8 | Stabilizing | -0.495 | Destabilizing | 1.62 | Stabilizing |
| 4 | -0.895 | Destabilizing | 0.57 | Stabilizing | -0.246 | Destabilizing | 3.15 | Stabilizing |
| 5 | 0.272 | Stabilizing | -0.13 | Destabilizing | 0.501 | Stabilizing | -0.3 | Destabilizing |
| 6 | -0.093 | Destabilizing | 0.1 | Stabilizing | -0.239 | Destabilizing | -0.39 | Destabilizing |
| 7 | -0.167 | Destabilizing | -0.83 | Destabilizing | -0.264 | Destabilizing | -1.33 | Destabilizing |
| 8 | 0.012 | Stabilizing | 0.12 | Stabilizing | 0.216 | Stabilizing | 3.89 | Stabilizing |
| 9 | -0.91 | Destabilizing | -1.48 | Destabilizing | -0.933 | Destabilizing | 1.32 | Stabilizing |
| 10 | -0.438 | Destabilizing | -0.34 | Destabilizing | -0.104 | Destabilizing | 2.86 | Stabilizing |
| 11 | -0.606 | Destabilizing | -2.2 | Destabilizing | -0.693 | Destabilizing | 1.71 | Stabilizing |
| 12 | -0.362 | Destabilizing | 0.05 | Stabilizing | -0.011 | Destabilizing | 0.52 | Stabilizing |
| 13 | -0.084 | Destabilizing | 1.97 | Stabilizing | 0.712 | Stabilizing | 3.47 | Stabilizing |
| 14 | -1.41 | Destabilizing | -4.41 | Destabilizing | -2.096 | Destabilizing | 1.7 | Stabilizing |
| 15 | -1.2 | Destabilizing | 0.02 | Stabilizing | -0.629 | Destabilizing | 2.48 | Stabilizing |
| 16 | -0.335 | Destabilizing | 0.84 | Stabilizing | 0.075 | Stabilizing | 1.24 | Stabilizing |
| 17 | -0.067 | Destabilizing | -0.05 | Destabilizing | 0.059 | Stabilizing | 0.37 | Stabilizing |
| 18 | 0.592 | Stabilizing | 0.46 | Stabilizing | 0.756 | Stabilizing | 0.09 | Stabilizing |
| 19 | -0.461 | Destabilizing | 0.03 | Stabilizing | -0.315 | Destabilizing | 1.65 | Stabilizing |
| 20 | -0.918 | Destabilizing | 0.45 | Stabilizing | -0.419 | Destabilizing | 0.88 | Stabilizing |
| 21 | -0.93 | Destabilizing | -0.18 | Destabilizing | -0.639 | Destabilizing | 1.17 | Stabilizing |
| 22 | -1.59 | Destabilizing | -1.49 | Destabilizing | -1.815 | Destabilizing | -1.02 | Destabilizing |
| 23 | -0.285 | Destabilizing | -0.73 | Destabilizing | -0.208 | Destabilizing | 1.3 | Stabilizing |
| 24 | -1.306 | Destabilizing | -0.11 | Destabilizing | -1.222 | Destabilizing | 1.24 | Stabilizing |
| 25 | -2.117 | Destabilizing | -0.96 | Destabilizing | -2.096 | Destabilizing | 0.06 | Stabilizing |
| 26 | -1.033 | Destabilizing | -0.39 | Destabilizing | -0.988 | Destabilizing | 1.02 | Stabilizing |
| 27 | 0.244 | Stabilizing | 0.82 | Stabilizing | 0.613 | Stabilizing | 1.36 | Stabilizing |
| 28 | -0.362 | Destabilizing | 0.23 | Stabilizing | -0.09 | Destabilizing | 0.98 | Stabilizing |
| 29 | -0.727 | Destabilizing | -1.23 | Destabilizing | -0.767 | Destabilizing | 0.24 | Stabilizing |
| 30 | -0.444 | Destabilizing | 1.29 | Stabilizing | 0.15 | Stabilizing | 0.98 | Stabilizing |
| 31 | -0.757 | Destabilizing | 0.69 | Stabilizing | -0.399 | Destabilizing | 0.67 | Stabilizing |
| 32 | -1.466 | Destabilizing | -1.98 | Destabilizing | -1.858 | Destabilizing | -0.01 | Destabilizing |
| 33 | -1.788 | Destabilizing | -1.76 | Destabilizing | -2.092 | Destabilizing | -1.13 | Destabilizing |
| 34 | -1.719 | Destabilizing | -0.63 | Destabilizing | -1.799 | Destabilizing | -0.41 | Destabilizing |
| 35 | -0.813 | Destabilizing | -0.19 | Destabilizing | -0.594 | Destabilizing | 0.31 | Stabilizing |
| 36 | -1.158 | Destabilizing | 0.16 | Stabilizing | -0.933 | Destabilizing | 0.56 | Stabilizing |
| 37 | -0.629 | Destabilizing | 0.62 | Stabilizing | -0.37 | Destabilizing | -0.33 | Destabilizing |
| 38 | 0.031 | Stabilizing | -0.39 | Destabilizing | 0.265 | Stabilizing | 0.69 | Stabilizing |
| 39 | -1.281 | Destabilizing | -1.19 | Destabilizing | -1.099 | Destabilizing | 1.2 | Stabilizing |
| 40 | 0.221 | Stabilizing | -0.46 | Destabilizing | 0.331 | Stabilizing | 1.18 | Stabilizing |
| 41 | -1.134 | Destabilizing | -0.41 | Destabilizing | -0.914 | Destabilizing | 2.25 | Stabilizing |
| 42 | -1.206 | Destabilizing | -0.15 | Destabilizing | -0.975 | Destabilizing | 1.94 | Stabilizing |
| 43 | -0.432 | Destabilizing | 0.61 | Stabilizing | 0.112 | Stabilizing | 1.53 | Stabilizing |
| 44 | -1.133 | Destabilizing | -0.72 | Destabilizing | -1.033 | Destabilizing | -0.49 | Destabilizing |
| 45 | -1.928 | Destabilizing | -1.39 | Destabilizing | -2.109 | Destabilizing | -0.97 | Destabilizing |
| 46 | -1.616 | Destabilizing | -0.89 | Destabilizing | -1.57 | Destabilizing | -1.36 | Destabilizing |
| 47 | -0.686 | Destabilizing | -1.46 | Destabilizing | -0.677 | Destabilizing | 2.23 | Stabilizing |
| 48 | -0.992 | Destabilizing | -0.72 | Destabilizing | -1.05 | Destabilizing | 0.62 | Stabilizing |
| 49 | 0.253 | Stabilizing | -0.4 | Destabilizing | 0.295 | Stabilizing | 0.06 | Stabilizing |
| 51 | -0.58 | Destabilizing | -0.7 | Destabilizing | -0.51 | Destabilizing | 1.12 | Stabilizing |
| 52 | -0.986 | Destabilizing | 0.18 | Stabilizing | -0.753 | Destabilizing | 0.21 | Stabilizing |
| 53 | -0.145 | Destabilizing | 0.4 | Stabilizing | 0.156 | Stabilizing | 0.48 | Stabilizing |
| 54 | -1.161 | Destabilizing | -0.69 | Destabilizing | -1.01 | Destabilizing | -0.18 | Destabilizing |
| 55 | -0.331 | Destabilizing | -0.35 | Destabilizing | -0.322 | Destabilizing | 2.16 | Stabilizing |
| 56 | -1.089 | Destabilizing | -1.39 | Destabilizing | -1.376 | Destabilizing | 3.43 | Stabilizing |
| 57 | -1.241 | Destabilizing | -3.02 | Destabilizing | -1.344 | Destabilizing | 1.82 | Stabilizing |
| 58 | -0.739 | Destabilizing | -1.97 | Destabilizing | -0.738 | Destabilizing | 1.03 | Stabilizing |
| 59 | -1.167 | Destabilizing | -0.95 | Destabilizing | -1.224 | Destabilizing | 2.06 | Stabilizing |
| 60 | -0.498 | Destabilizing | 1.42 | Stabilizing | 0.222 | Stabilizing | 1.29 | Stabilizing |
| 60 | -0.556 | Destabilizing | -0.14 | Destabilizing | -0.198 | Destabilizing | 3 | Stabilizing |
| 61 | -0.056 | Destabilizing | -1.4 | Destabilizing | -0.171 | Destabilizing | -1.41 | Destabilizing |
| 62 | -0.243 | Destabilizing | -0.53 | Destabilizing | -0.242 | Destabilizing | -1.75 | Destabilizing |
| 63 | -0.676 | Destabilizing | -0.16 | Destabilizing | -0.612 | Destabilizing | -0.49 | Destabilizing |
| 64 | -0.239 | Destabilizing | 0.34 | Stabilizing | 0.161 | Stabilizing | 1.41 | Stabilizing |
| 65 | 0.465 | Stabilizing | 3.13 | Stabilizing | 1.363 | Stabilizing | 3.94 | Stabilizing |
| 66 | -0.87 | Destabilizing | -0.78 | Destabilizing | -1.011 | Destabilizing | 3.59 | Stabilizing |
| 67 | -0.035 | Destabilizing | -0.67 | Destabilizing | -0.006 | Destabilizing | 0.72 | Stabilizing |
| 68 | -0.372 | Destabilizing | -0.58 | Destabilizing | -0.176 | Destabilizing | 3.03 | Stabilizing |
| 69 | -0.577 | Destabilizing | -0.86 | Destabilizing | -0.496 | Destabilizing | 1.39 | Stabilizing |
| 70 | -1.772 | Destabilizing | -1.01 | Destabilizing | -1.725 | Destabilizing | 1.82 | Stabilizing |
| 71 | -0.883 | Destabilizing | -0.23 | Destabilizing | -0.524 | Destabilizing | 0.76 | Stabilizing |
| 72 | -0.669 | Destabilizing | -0.61 | Destabilizing | -0.439 | Destabilizing | 2.64 | Stabilizing |
| 73 | -0.443 | Destabilizing | -0.73 | Destabilizing | -0.388 | Destabilizing | 2.58 | Stabilizing |
| 74 | -0.772 | Destabilizing | -0.39 | Destabilizing | -0.495 | Destabilizing | 2.19 | Stabilizing |
| 75 | -0.675 | Destabilizing | 2.26 | Stabilizing | 0.211 | Stabilizing | 1.67 | Stabilizing |
| 76 | -0.66 | Destabilizing | -1.12 | Destabilizing | -0.6 | Destabilizing | 1.6 | Stabilizing |
| 77 | -0.146 | Destabilizing | -0.17 | Destabilizing | -0.11 | Destabilizing | -0.7 | Destabilizing |
| 78 | -1.025 | Destabilizing | 0.15 | Stabilizing | -0.864 | Destabilizing | -0.39 | Destabilizing |
| 79 | -0.469 | Destabilizing | -0.26 | Destabilizing | -0.288 | Destabilizing | 0.05 | Stabilizing |
| 80 | -0.599 | Destabilizing | 0.95 | Stabilizing | 0.036 | Stabilizing | 2.68 | Stabilizing |
| 81 | -0.326 | Destabilizing | -2.07 | Destabilizing | -0.598 | Destabilizing | 2.28 | Stabilizing |
| 82 | -0.06 | Destabilizing | 0.880 | Stabilizing | 0.495 | Stabilizing | 2.22 | Stabilizing |
| 83 | -0.466 | Destabilizing | -0.43 | Destabilizing | -0.336 | Destabilizing | 2.76 | Stabilizing |
| 84 | -1.287 | Destabilizing | -1.93 | Destabilizing | -1.314 | Destabilizing | 1.93 | Stabilizing |
| 85 | -0.596 | Destabilizing | -0.01 | Destabilizing | -0.315 | Destabilizing | 0.67 | Stabilizing |
| 86 | -0.962 | Destabilizing | -0.820 | Destabilizing | -0.584 | Destabilizing | 1.81 | Stabilizing |
| 87 | -0.835 | Destabilizing | -0.25 | Destabilizing | -0.518 | Destabilizing | 1.7 | Stabilizing |
| 88 | -0.298 | Destabilizing | -1.57 | Destabilizing | -0.587 | Destabilizing | 1.94 | Stabilizing |
| 89 | -0.677 | Destabilizing | -0.28 | Destabilizing | -0.57 | Destabilizing | -0.38 | Destabilizing |
| 90 | -1.431 | Destabilizing | 0.770 | Stabilizing | -0.898 | Destabilizing | -1.38 | Destabilizing |
| 91 | -0.693 | Destabilizing | -1.27 | Destabilizing | -0.759 | Destabilizing | -0.19 | Destabilizing |
| 92 | -1.337 | Destabilizing | -1.1 | Destabilizing | -1.139 | Destabilizing | 1.84 | Stabilizing |
| 93 | -0.233 | Destabilizing | 0.28 | Stabilizing | 0.047 | Stabilizing | -0.18 | Destabilizing |
| 94 | 0.416 | Stabilizing | -1.53 | Destabilizing | 0.324 | Stabilizing | -0.54 | Destabilizing |
| 95 | -1.163 | Destabilizing | -1.73 | Destabilizing | -1.367 | Destabilizing | -0.35 | Destabilizing |
| 96 | 0.074 | Stabilizing | 0.11 | Stabilizing | 0.421 | Stabilizing | 1.25 | Stabilizing |
| 97 | -1.255 | Destabilizing | -0.79 | Destabilizing | -1.177 | Destabilizing | -1.41 | Destabilizing |
| 98 | -0.201 | Destabilizing | -0.201 | Destabilizing | -0.434 | Destabilizing | -0.37 | Destabilizing |
| 99 | -0.296 | Destabilizing | -0.14 | Destabilizing | -0.004 | Destabilizing | 2.14 | Stabilizing |
| 100 | -0.155 | Destabilizing | -0.1 | Destabilizing | 0.045 | Stabilizing | -0.56 | Destabilizing |
| 101 | 0.116 | Stabilizing | -0.75 | Destabilizing | 0.267 | Stabilizing | 0.93 | Stabilizing |
| 102 | -1.023 | Destabilizing | -1.53 | Destabilizing | -1.087 | Destabilizing | 1.72 | Stabilizing |
| 103 | 0.043 | Stabilizing | -0.58 | Destabilizing | -0.035 | Destabilizing | 1.74 | Stabilizing |
| 104 | -0.143 | Destabilizing | -0.24 | Destabilizing | 0.07 | Stabilizing | 0.66 | Stabilizing |
| 105 | 0.125 | Stabilizing | 0.78 | Stabilizing | 0.409 | Stabilizing | 0.75 | Stabilizing |
| 106 | -0.822 | Destabilizing | 1.030 | Stabilizing | -0.194 | Destabilizing | 2.72 | Stabilizing |
| 107 | 0.007 | Stabilizing | 0.69 | Stabilizing | 0.607 | Stabilizing | 0.97 | Stabilizing |
| 108 | -0.126 | Destabilizing | -0.49 | Destabilizing | 0.205 | Stabilizing | 0.45 | Stabilizing |
| 109 | -0.617 | Destabilizing | -0.31 | Destabilizing | -0.467 | Destabilizing | 2.02 | Stabilizing |
| 110 | -1.082 | Destabilizing | -0.39 | Destabilizing | -1.089 | Destabilizing | 0.01 | Stabilizing |
| 111 | 0.075 | Stabilizing | 1.3 | Stabilizing | 0.576 | Stabilizing | 2.53 | Stabilizing |
| 112 | -0.908 | Destabilizing | -1.4 | Destabilizing | -1.038 | Destabilizing | 0.58 | Stabilizing |
| 113 | -0.264 | Destabilizing | 0.100 | Stabilizing | -0.355 | Destabilizing | -0.12 | Destabilizing |
| 114 | -1.842 | Destabilizing | -0.37 | Destabilizing | -2.011 | Destabilizing | -0.06 | Destabilizing |
| 115 | -1.052 | Destabilizing | 0 | Stabilizing | -0.927 | Destabilizing | 1.44 | Stabilizing |
| 116 | -0.404 | Destabilizing | 3.38 | Stabilizing | 0.629 | Stabilizing | 2.62 | Stabilizing |
| 117 | -2.477 | Destabilizing | -0.71 | Destabilizing | -2.31 | Destabilizing | 1.65 | Stabilizing |
| 118 | -0.113 | Destabilizing | 1.28 | Stabilizing | 0.536 | Stabilizing | 3.15 | Stabilizing |
| 119 | -1.829 | Destabilizing | -1.17 | Destabilizing | -1.856 | Destabilizing | 2.81 | Stabilizing |
| 120 | -3.028 | Destabilizing | -3.48 | Destabilizing | -3.24 | Destabilizing | 0.55 | Stabilizing |
| 121 | -0.873 | Destabilizing | -2.63 | Destabilizing | -1.173 | Destabilizing | 0.94 | Stabilizing |
| 122 | -0.277 | Destabilizing | 0.36 | Stabilizing | 0.216 | Stabilizing | 2.47 | Stabilizing |
| 123 | -0.746 | Destabilizing | -0.8 | Destabilizing | -0.694 | Destabilizing | 0.91 | Stabilizing |
| 124 | -1.079 | Destabilizing | 0.81 | Stabilizing | -0.623 | Destabilizing | -0.53 | Destabilizing |
| 125 | -0.697 | Destabilizing | 0.370 | Stabilizing | -0.202 | Destabilizing | -0.27 | Destabilizing |
| 126 | -1.189 | Destabilizing | -0.75 | Destabilizing | -1.039 | Destabilizing | 1.16 | Stabilizing |
| 127 | -1.101 | Destabilizing | 0.15 | Stabilizing | -1.023 | Destabilizing | 0.59 | Stabilizing |
| 128 | -0.412 | Destabilizing | 0.34 | Stabilizing | -0.243 | Destabilizing | 1.68 | Stabilizing |
| 129 | -0.268 | Destabilizing | 0.82 | Stabilizing | 0.248 | Stabilizing | 0.54 | Stabilizing |
| 130 | -0.326 | Destabilizing | 0.05 | Stabilizing | 0.044 | Stabilizing | 0.56 | Stabilizing |
| 131 | -0.661 | Destabilizing | -0.52 | Destabilizing | -0.405 | Destabilizing | 1.06 | Stabilizing |
| 132 | -0.538 | Destabilizing | -0.21 | Destabilizing | -0.276 | Destabilizing | 2.76 | Stabilizing |
| 133 | -0.47 | Destabilizing | -0.32 | Destabilizing | -0.043 | Destabilizing | 1.44 | Stabilizing |
| 134 | -0.787 | Destabilizing | 0.04 | Stabilizing | -0.617 | Destabilizing | 0.68 | Stabilizing |
| 135 | -1.078 | Destabilizing | -2.32 | Destabilizing | -1.169 | Destabilizing | 1.98 | Stabilizing |
| 136 | -0.114 | Destabilizing | 2.26 | Stabilizing | 0.74 | Stabilizing | 2.96 | Stabilizing |
| 137 | -1.256 | Destabilizing | -2.85 | Destabilizing | -1.642 | Destabilizing | 1.27 | Stabilizing |
| 138 | -0.125 | Destabilizing | 1.26 | Stabilizing | 0.408 | Stabilizing | 2.14 | Stabilizing |
| 139 | -0.335 | Destabilizing | -0.270 | Destabilizing | -0.364 | Destabilizing | -0.98 | Destabilizing |
| 140 | -0.706 | Destabilizing | 0.26 | Stabilizing | -0.638 | Destabilizing | -0.4 | Destabilizing |
| 141 | 0.302 | Stabilizing | -0.46 | Destabilizing | 0.213 | Stabilizing | -1.47 | Destabilizing |
| 142 | -0.405 | Destabilizing | 0.19 | Stabilizing | -0.241 | Destabilizing | 2.88 | Stabilizing |
| 143 | -0.815 | Destabilizing | 1.17 | Stabilizing | -0.138 | Destabilizing | 2.98 | Stabilizing |
| 144 | 0.346 | Stabilizing | 0.28 | Stabilizing | 0.617 | Stabilizing | 1.71 | Stabilizing |
| 145 | -1.334 | Destabilizing | -2.38 | Destabilizing | -1.62 | Destabilizing | 1.42 | Stabilizing |
| 146 | -1.196 | Destabilizing | -2.82 | Destabilizing | -1.752 | Destabilizing | -0.46 | Destabilizing |
| 147 | -0.091 | Destabilizing | -0.52 | Destabilizing | -0.075 | Destabilizing | 2.58 | Stabilizing |
| 148 | -0.269 | Destabilizing | -0.5 | Destabilizing | -0.25 | Destabilizing | -0.28 | Destabilizing |
| 149 | -0.551 | Destabilizing | -0.48 | Destabilizing | -0.442 | Destabilizing | 0.61 | Stabilizing |
| 150 | -1.854 | Destabilizing | -2.9 | Destabilizing | -2.096 | Destabilizing | -0.24 | Destabilizing |
| 151 | -0.827 | Destabilizing | 2.630 | Stabilizing | 0.138 | Stabilizing | 2.35 | Stabilizing |
| 152 | -0.054 | Destabilizing | -0.57 | Destabilizing | -0.008 | Destabilizing | -0.38 | Destabilizing |
| 153 | -2.282 | Destabilizing | -1.57 | Destabilizing | -2.327 | Destabilizing | 1.11 | Stabilizing |
| 154 | -0.496 | Destabilizing | -0.28 | Destabilizing | -0.151 | Destabilizing | 1.68 | Stabilizing |
| 155 | -0.424 | Destabilizing | -1.24 | Destabilizing | -0.319 | Destabilizing | 1.1 | Stabilizing |
| 156 | -0.725 | Destabilizing | 0.25 | Stabilizing | -0.482 | Destabilizing | 2.54 | Stabilizing |
| 157 | -0.711 | Destabilizing | 0.31 | Stabilizing | -0.2 | Destabilizing | 1.55 | Stabilizing |
| 158 | -0.58 | Destabilizing | -0.61 | Destabilizing | -0.206 | Destabilizing | 0.7 | Stabilizing |
| 159 | -0.484 | Destabilizing | 0.33 | Stabilizing | -0.079 | Destabilizing | -1.15 | Destabilizing |
| 160 | -0.904 | Destabilizing | -3.12 | Destabilizing | -1.094 | Destabilizing | 2.93 | Stabilizing |
| 161 | 0.195 | Stabilizing | 0.1 | Stabilizing | -0.043 | Destabilizing | -0.1 | Destabilizing |
| 162 | -1.619 | Destabilizing | -0.92 | Destabilizing | -1.648 | Destabilizing | 1.68 | Stabilizing |
| 163 | -2.073 | Destabilizing | -1.82 | Destabilizing | -2.074 | Destabilizing | 1.95 | Stabilizing |
| 164 | -1.074 | Destabilizing | 0.78 | Stabilizing | -0.363 | Destabilizing | 3.12 | Stabilizing |
| 165 | -1.092 | Destabilizing | -0.86 | Destabilizing | -1.149 | Destabilizing | 3.59 | Stabilizing |
| 166 | -0.247 | Destabilizing | -0.24 | Destabilizing | -0.222 | Destabilizing | -1.75 | Destabilizing |
| 167 | -0.244 | Destabilizing | -1.000 | Destabilizing | -0.288 | Destabilizing | -1.89 | Destabilizing |
| 168 | 0.277 | Stabilizing | 0.2 | Stabilizing | 0.418 | Stabilizing | -1.05 | Destabilizing |
| 169 | 0.05 | Stabilizing | -0.81 | Destabilizing | -0.066 | Destabilizing | -0.26 | Destabilizing |
| 170 | -0.642 | Destabilizing | -0.79 | Destabilizing | -0.525 | Destabilizing | 1.09 | Stabilizing |
| 171 | -0.805 | Destabilizing | -0.97 | Destabilizing | -0.65 | Destabilizing | 1.36 | Stabilizing |
| 172 | -0.882 | Destabilizing | -2.54 | Destabilizing | -1.2 | Destabilizing | 1.89 | Stabilizing |
| 173 | -0.69 | Destabilizing | 0.05 | Stabilizing | -0.646 | Destabilizing | 1.51 | Stabilizing |
| 174 | -0.332 | Destabilizing | 0.32 | Stabilizing | -0.102 | Destabilizing | 1.59 | Stabilizing |
| 175 | -0.335 | Destabilizing | 0.31 | Stabilizing | -0.401 | Destabilizing | -0.62 | Destabilizing |
| 176 | 0.105 | Stabilizing | -0.11 | Destabilizing | 0.399 | Stabilizing | 0.31 | Stabilizing |
| 177 | -0.762 | Destabilizing | -0.88 | Destabilizing | -0.534 | Destabilizing | 1.62 | Stabilizing |
| 178 | -0.669 | Destabilizing | -0.43 | Destabilizing | -0.554 | Destabilizing | 2.64 | Stabilizing |
| 179 | -0.521 | Destabilizing | 0.28 | Stabilizing | -0.575 | Destabilizing | -0.48 | Destabilizing |
| 180 | -1.544 | Destabilizing | -1.39 | Destabilizing | -1.807 | Destabilizing | 2.19 | Stabilizing |
| 181 | 0.012 | Stabilizing | 0.47 | Stabilizing | 0.277 | Stabilizing | 2.79 | Stabilizing |
| 182 | -0.427 | Destabilizing | -0.44 | Destabilizing | -0.285 | Destabilizing | 2.48 | Stabilizing |
| 183 | -1.735 | Destabilizing | -2.1 | Destabilizing | -1.899 | Destabilizing | 2.16 | Stabilizing |
| 184 | -0.605 | Destabilizing | -2.39 | Destabilizing | -0.984 | Destabilizing | 1.35 | Stabilizing |
| 185 | -2.751 | Destabilizing | -3.72 | Destabilizing | -3.067 | Destabilizing | 0.1 | Stabilizing |
| 186 | -0.583 | Destabilizing | -0.15 | Destabilizing | -0.341 | Destabilizing | 0.92 | Stabilizing |
| 187 | -0.029 | Destabilizing | -1.61 | Destabilizing | -0.163 | Destabilizing | 0.86 | Stabilizing |
| 188 | -0.762 | Destabilizing | 0.18 | Stabilizing | -0.199 | Destabilizing | 2.33 | Stabilizing |
| 189 | 0.007 | Stabilizing | -0.46 | Destabilizing | 0.325 | Stabilizing | 1.49 | Stabilizing |
| 190 | -0.21 | Destabilizing | -0.69 | Destabilizing | 0.011 | Stabilizing | 1.81 | Stabilizing |
| 191 | -1.033 | Destabilizing | 0.62 | Stabilizing | -0.584 | Destabilizing | 1.54 | Stabilizing |
| 192 | -0.545 | Destabilizing | -0.8 | Destabilizing | -0.423 | Destabilizing | 2.46 | Stabilizing |
| 193 | -0.014 | Destabilizing | 0.140 | Stabilizing | 0.361 | Stabilizing | 0.69 | Stabilizing |
| 194 | -0.956 | Destabilizing | -0.45 | Destabilizing | -0.972 | Destabilizing | 2.24 | Stabilizing |
| 195 | -0.136 | Destabilizing | 2.51 | Stabilizing | 0.534 | Stabilizing | -0.63 | Destabilizing |
| 196 | -0.46 | Destabilizing | 0.19 | Stabilizing | -0.016 | Destabilizing | 2.16 | Stabilizing |
| 197 | -1.733 | Destabilizing | -3.12 | Destabilizing | -2.005 | Destabilizing | 1.6 | Stabilizing |
| 198 | -1.085 | Destabilizing | -2.87 | Destabilizing | -1.46 | Destabilizing | -1.43 | Destabilizing |
| 199 | -0.006 | Destabilizing | -1.591 | Destabilizing | -0.111 | Stabilizing | 1.42 | Stabilizing |
| 200 | -0.78 | Destabilizing | -0.01 | Destabilizing | -0.539 | Destabilizing | 1.78 | Stabilizing |
| 201 | -0.1 | Destabilizing | -0.17 | Destabilizing | 0.210 | Stabilizing | 0.98 | Stabilizing |
| 202 | -2.028 | Destabilizing | -1.09 | Destabilizing | -2.16 | Destabilizing | -1.52 | Destabilizing |
| 203 | -2.193 | Destabilizing | -0.65 | Destabilizing | -2.271 | Destabilizing | -0.78 | Destabilizing |
| 204 | -1.729 | Destabilizing | -1.85 | Destabilizing | -2.095 | Destabilizing | -2.07 | Destabilizing |
| 205 | -1.016 | Destabilizing | -1.05 | Destabilizing | -0.879 | Destabilizing | 2.69 | Stabilizing |
| 206 | -1.769 | Destabilizing | -0.83 | Destabilizing | -1.981 | Destabilizing | 2.32 | Stabilizing |
| 207 | -0.62 | Destabilizing | -0.35 | Destabilizing | -0.296 | Destabilizing | 1.48 | Stabilizing |
| 208 | -0.471 | Destabilizing | -1.08 | Destabilizing | -0.404 | Destabilizing | 1.76 | Stabilizing |
| 209 | -0.877 | Destabilizing | -0.76 | Destabilizing | -0.932 | Destabilizing | 1.74 | Stabilizing |
| 210 | -0.486 | Destabilizing | 0.22 | Stabilizing | -0.106 | Destabilizing | 1.56 | Stabilizing |
| 211 | -1.274 | Destabilizing | -0.2 | Destabilizing | -1.204 | Destabilizing | 1.09 | Stabilizing |
| 212 | -0.579 | Destabilizing | -1 | Destabilizing | -0.77 | Destabilizing | -0.56 | Destabilizing |
| 213 | -0.903 | Destabilizing | -0.72 | Destabilizing | -0.89 | Destabilizing | 0.67 | Stabilizing |
| 214 | -0.131 | Destabilizing | 0.08 | Stabilizing | -0.312 | Destabilizing | 1.26 | Stabilizing |
| 215 | -0.297 | Destabilizing | -0.6 | Destabilizing | -0.017 | Destabilizing | 3.01 | Stabilizing |
| 216 | -1.334 | Destabilizing | -2.38 | Destabilizing | -1.62 | Destabilizing | 1.41 | Stabilizing |
| 217 | 0.904 | Stabilizing | -1.88 | Destabilizing | -1.62 | Destabilizing | 1.83 | Stabilizing |
| 218 | -0.575 | Destabilizing | -0.42 | Destabilizing | -0.365 | Destabilizing | 2.65 | Stabilizing |
| 219 | -0.297 | Destabilizing | 0.13 | Stabilizing | 0.055 | Stabilizing | 1.97 | Stabilizing |
| 220 | -0.101 | Destabilizing | 0.86 | Stabilizing | 0.34 | Stabilizing | 1.97 | Stabilizing |
| 221 | -0.521 | Destabilizing | 0.6 | Stabilizing | -0.37 | Destabilizing | 0.76 | Stabilizing |
| 222 | -1.464 | Destabilizing | 0.57 | Stabilizing | -1.285 | Destabilizing | -0.12 | Destabilizing |
| 223 | 0.157 | Stabilizing | 0.58 | Stabilizing | 0.324 | Stabilizing | -0.86 | Destabilizing |
| 224 | -0.135 | Destabilizing | -1.21 | Destabilizing | -0.476 | Destabilizing | -1.47 | Destabilizing |
| 225 | -0.907 | Destabilizing | -0.56 | Destabilizing | -0.75 | Destabilizing | 2.18 | Stabilizing |
| 226 | -0.086 | Destabilizing | 0.57 | Stabilizing | 0.428 | Stabilizing | 0.39 | Stabilizing |
| 227 | -0.522 | Destabilizing | -0.97 | Destabilizing | -0.372 | Destabilizing | 2.35 | Stabilizing |
| 228 | -0.256 | Destabilizing | 2.43 | Stabilizing | 0.603 | Stabilizing | 0.73 | Stabilizing |
| 229 | -1.447 | Destabilizing | -1.53 | Destabilizing | -1.551 | Destabilizing | 2.71 | Stabilizing |
| 230 | -1.5 | Destabilizing | -0.74 | Destabilizing | -1.625 | Destabilizing | 0.43 | Stabilizing |
| 231 | -1.405 | Destabilizing | -1.76 | Destabilizing | -1.721 | Destabilizing | -1 | Destabilizing |
| 232 | 0.839 | Stabilizing | 0.25 | Stabilizing | 1.072 | Stabilizing | 0.92 | Stabilizing |
| 233 | -1.967 | Destabilizing | -3.05 | Destabilizing | -2.237 | Destabilizing | -0.21 | Destabilizing |
| 234 | -1.058 | Destabilizing | -1.76 | Destabilizing | -0.933 | Destabilizing | 0.47 | Stabilizing |
| 235 | -1.384 | Destabilizing | -0.63 | Destabilizing | -1.257 | Destabilizing | -1.45 | Destabilizing |
| 236 | -1.445 | Destabilizing | 0.72 | Stabilizing | -0.979 | Destabilizing | -0.75 | Destabilizing |
| 237 | -0.231 | Destabilizing | 0.32 | Stabilizing | 0.004 | Stabilizing | 0.8 | Stabilizing |
| 238 | -0.048 | Destabilizing | 0.01 | Stabilizing | -0.042 | Destabilizing | 1.45 | Stabilizing |
| 239 | -1.747 | Destabilizing | -2.35 | Destabilizing | -1.849 | Destabilizing | 3.59 | Stabilizing |
| 240 | -1.437 | Destabilizing | 0.62 | Stabilizing | -0.787 | Destabilizing | 3.42 | Stabilizing |
| 241 | -1.845 | Destabilizing | 0.4 | Stabilizing | -1.282 | Destabilizing | 2.49 | Stabilizing |
| 242 | -2.251 | Destabilizing | -2.02 | Destabilizing | -2.142 | Destabilizing | 1.82 | Stabilizing |
| 243 | -2.015 | Destabilizing | -2.66 | Destabilizing | -2.436 | Destabilizing | 0.08 | Stabilizing |
| 244 | -0.525 | Destabilizing | -0.08 | Destabilizing | -0.319 | Destabilizing | 1.92 | Stabilizing |
| 245 | -0.347 | Destabilizing | -0.18 | Destabilizing | -0.222 | Destabilizing | -0.23 | Destabilizing |
| 246 | -0.434 | Destabilizing | -0.55 | Destabilizing | -0.445 | Destabilizing | 0.27 | Stabilizing |
| 247 | -0.255 | Destabilizing | 0.14 | Stabilizing | -0.435 | Destabilizing | 0.05 | Stabilizing |
| 248 | -1.735 | Destabilizing | -1.12 | Destabilizing | -1.75 | Destabilizing | 2.53 | Stabilizing |
| 249 | -1.265 | Destabilizing | -1.53 | Destabilizing | -1.358 | Destabilizing | 3.23 | Stabilizing |
| 250 | -0.781 | Destabilizing | -0.39 | Destabilizing | -0.795 | Destabilizing | -0.26 | Destabilizing |
| 251 | 0.104 | Stabilizing | -0.61 | Destabilizing | 0.051 | Stabilizing | -1.75 | Destabilizing |
| 252 | 0.468 | Stabilizing | -1.65 | Destabilizing | 0.091 | Stabilizing | -2.3 | Destabilizing |
| 253 | 0.058 | Stabilizing | -1.65 | Destabilizing | -0.277 | Destabilizing | -1.26 | Destabilizing |
| 254 | -0.196 | Destabilizing | -0.87 | Destabilizing | -0.145 | Destabilizing | -0.2 | Destabilizing |
| 255 | -0.183 | Destabilizing | 0.04 | Stabilizing | -0.311 | Destabilizing | -0.67 | Destabilizing |
| 256 | -0.883 | Destabilizing | -0.66 | Destabilizing | -0.81 | Destabilizing | 1.17 | Stabilizing |
| 257 | -0.679 | Destabilizing | -0.34 | Destabilizing | -0.530 | Destabilizing | 1.74 | Stabilizing |
| 258 | -1.459 | Destabilizing | 0.35 | Stabilizing | -1.337 | Destabilizing | 0.2 | Stabilizing |
| 259 | -0.034 | Destabilizing | -1 | Destabilizing | -0.263 | Destabilizing | -0.99 | Destabilizing |
| 260 | -0.782 | Destabilizing | -1.18 | Destabilizing | -0.95 | Destabilizing | -0.92 | Destabilizing |
| 261 | 0.859 | Stabilizing | -0.06 | Destabilizing | 1.013 | Stabilizing | 0.87 | Stabilizing |
| 262 | -0.166 | Destabilizing | 1.93 | Stabilizing | 0.584 | Stabilizing | 2.23 | Stabilizing |
| 263 | -0.285 | Destabilizing | 0.39 | Stabilizing | 0.125 | Stabilizing | 1.88 | Stabilizing |
| 264 | -0.607 | Destabilizing | -0.01 | Destabilizing | -0.288 | Destabilizing | 1.93 | Stabilizing |
| 265 | -0.037 | Destabilizing | 0.48 | Stabilizing | 0.304 | Stabilizing | 2.76 | Stabilizing |
| 266 | -0.411 | Destabilizing | -0.94 | Destabilizing | -0.322 | Destabilizing | 1.28 | Stabilizing |
| 267 | -0.513 | Destabilizing | -0.57 | Destabilizing | -0.334 | Destabilizing | 0.34 | Stabilizing |
| 268 | -0.059 | Destabilizing | 0.49 | Stabilizing | 0.293 | Stabilizing | 1.58 | Stabilizing |
| 269 | -0.661 | Destabilizing | -0.35 | Destabilizing | -0.59 | Destabilizing | 2.23 | Stabilizing |
| 270 | -0.083 | Destabilizing | -0.16 | Destabilizing | 0.064 | Stabilizing | 0.4 | Stabilizing |
| 271 | -0.119 | Destabilizing | 0.360 | Stabilizing | -0.073 | Destabilizing | 0.47 | Stabilizing |
| 272 | -1.214 | Destabilizing | 1.62 | Stabilizing | -0.628 | Destabilizing | 2.38 | Stabilizing |
| 273 | -0.885 | Destabilizing | -1.58 | Destabilizing | -0.927 | Destabilizing | 1.79 | Stabilizing |
| 274 | 0.183 | Stabilizing | -1.65 | Destabilizing | -0.168 | Destabilizing | -1.87 | Destabilizing |
| 275 | -0.106 | Destabilizing | -1.28 | Destabilizing | 0.202 | Stabilizing | 1.14 | Stabilizing |
| 276 | -0.304 | Destabilizing | -1.46 | Destabilizing | -0.261 | Destabilizing | 1.72 | Stabilizing |

### Cancer-causing mutation analysis

A set of 64 mutations commonly identified by different algorithms was subjected to FATHMM server to determine mutations that could have a potential role in cancer-causing/cancer-promoting. Seven mutations such as G47R, K48R, R75I, R324H, D681N, R683Q, and R683W were predicted to be cancer-causing/cancer-promoting [Table 4] and were residing on different domains of *XPD.*

**Table 4.**
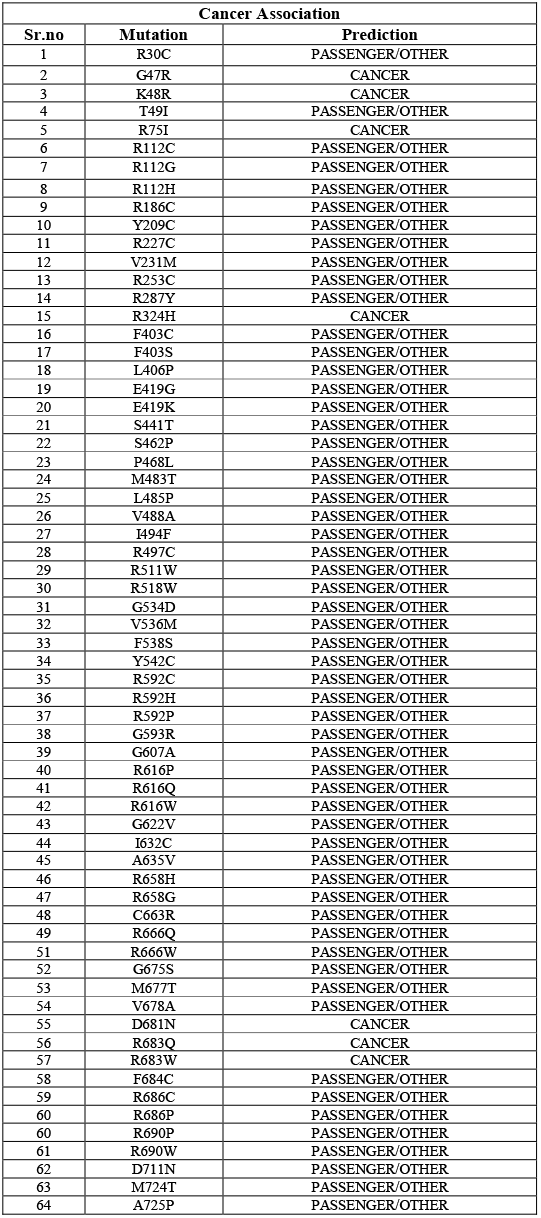
Prediction Cancer association using FATHMM Server

| Cancer Association |  |  |
| --- | --- | --- |
| Sr.no | Mutation | Prediction |
| 1 | R30C | PASSENGER/OTHER |
| 2 | G47R | CANCER |
| 3 | K48R | CANCER |
| 4 | T49I | PASSENGER/OTHER |
| 5 | R75I | CANCER |
| 6 | R112C | PASSENGER/OTHER |
| 7 | R112G | PASSENGER/OTHER |
| 8 | R112H | PASSENGER/OTHER |
| 9 | R186C | PASSENGER/OTHER |
| 10 | Y209C | PASSENGER/OTHER |
| 11 | R227C | PASSENGER/OTHER |
| 12 | V231M | PASSENGER/OTHER |
| 13 | R253C | PASSENGER/OTHER |
| 14 | R287Y | PASSENGER/OTHER |
| 15 | R324H | CANCER |
| 16 | F403C | PASSENGER/OTHER |
| 17 | F403S | PASSENGER/OTHER |
| 18 | L406P | PASSENGER/OTHER |
| 19 | E419G | PASSENGER/OTHER |
| 20 | E419K | PASSENGER/OTHER |
| 21 | S441T | PASSENGER/OTHER |
| 22 | S462P | PASSENGER/OTHER |
| 23 | P468L | PASSENGER/OTHER |
| 24 | M483T | PASSENGER/OTHER |
| 25 | L485P | PASSENGER/OTHER |
| 26 | V488A | PASSENGER/OTHER |
| 27 | I494F | PASSENGER/OTHER |
| 28 | R497C | PASSENGER/OTHER |
| 29 | R511W | PASSENGER/OTHER |
| 30 | R518W | PASSENGER/OTHER |
| 31 | G534D | PASSENGER/OTHER |
| 32 | V536M | PASSENGER/OTHER |
| 33 | F538S | PASSENGER/OTHER |
| 34 | Y542C | PASSENGER/OTHER |
| 35 | R592C | PASSENGER/OTHER |
| 36 | R592H | PASSENGER/OTHER |
| 37 | R592P | PASSENGER/OTHER |
| 38 | G593R | PASSENGER/OTHER |
| 39 | G607A | PASSENGER/OTHER |
| 40 | R616P | PASSENGER/OTHER |
| 41 | R616Q | PASSENGER/OTHER |
| 42 | R616W | PASSENGER/OTHER |
| 43 | G622V | PASSENGER/OTHER |
| 44 | I632C | PASSENGER/OTHER |
| 45 | A635V | PASSENGER/OTHER |
| 46 | R658H | PASSENGER/OTHER |
| 47 | R658G | PASSENGER/OTHER |
| 48 | C663R | PASSENGER/OTHER |
| 49 | R666Q | PASSENGER/OTHER |
| 51 | R666W | PASSENGER/OTHER |
| 52 | G675S | PASSENGER/OTHER |
| 53 | M677T | PASSENGER/OTHER |
| 54 | V678A | PASSENGER/OTHER |
| 55 | D681N | CANCER |
| 56 | R683Q | CANCER |
| 57 | R683W | CANCER |
| 58 | F684C | PASSENGER/OTHER |
| 59 | R686C | PASSENGER/OTHER |
| 60 | R686P | PASSENGER/OTHER |
| 60 | R690P | PASSENGER/OTHER |
| 61 | R690W | PASSENGER/OTHER |
| 62 | D711N | PASSENGER/OTHER |
| 63 | M724T | PASSENGER/OTHER |
| 64 | A725P | PASSENGER/OTHER |

### Conservation analysis

Of the 7 cancer-causing mutations, K48R was taken for further comprehensive analysis and subjected to conservation analysis using the ConSurf server tool. The result obtained revealed that the K48 position was conserved at a scale of 6 [Figure 2]. Based on this analysis the K48R mutation was chosen for subsequent structural analysis in a computational simulated environment to determine its impact on protein stability and function.

**Figure 2.**
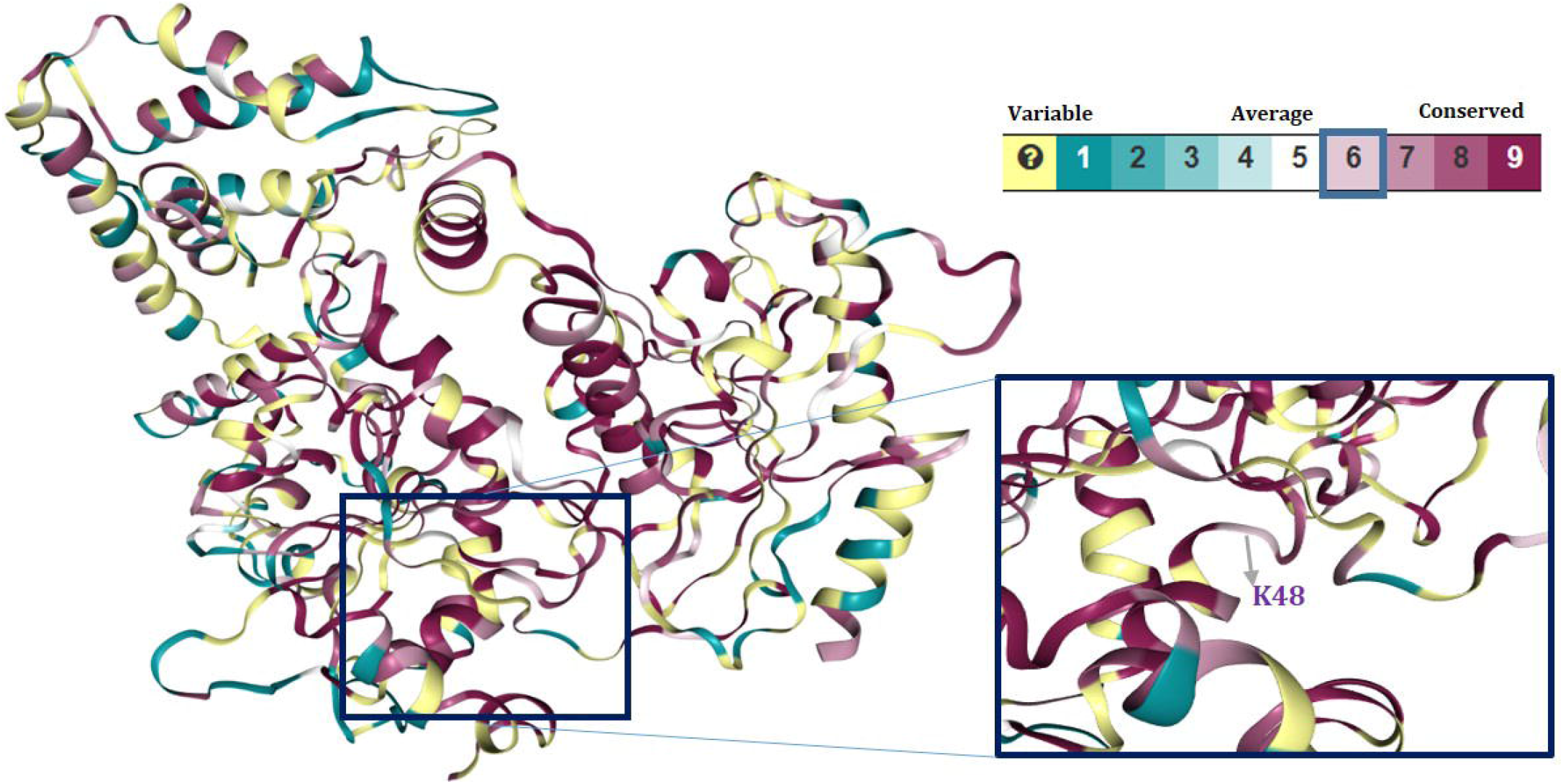
Residue conservation analysis of K48th residue of XPD using ConSurf Server

### Construction of a mutant model

Crystal structure with PDB id 5IVW was taken as a native control model for the analysis. For constructing the mutant model of K48R, a mutant sequence was submitted to I-TASSER server which first retrieved template protein of similar folds from the PDB library by locally installed Meta threading approach, where it generated ten thousand conformations (decoys). Then SPICKER program was used by I-TASSER to cluster all the decoys based on pairwise structure alignment and report up to five models that correspond to the five largest clusters. Model 1 with the best confidence score (C-score) [Supplementary Table 5] was taken for further analysis [Figure 3A, 3B].

**Figure 3.**
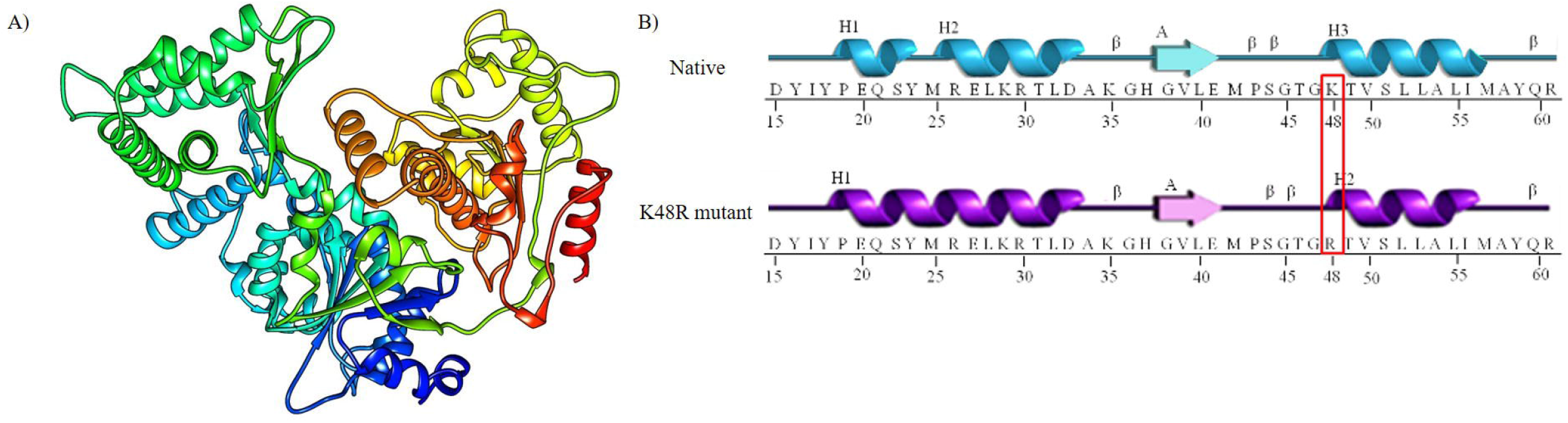
3D mutant model **A:** 3D Mutant Model (K48R) by I-TASSER server **B:** Substitution of Arginine for Lysine at 48^th^ residue of XPD

Quality assessment of the model 1 was done using Ramachandran plot where 86.8% of the residues were found to be in the favoured region and analogous to Ramachandran plot derived for native XPD structure [Figure 4A, 4B].

**Figure 4.**
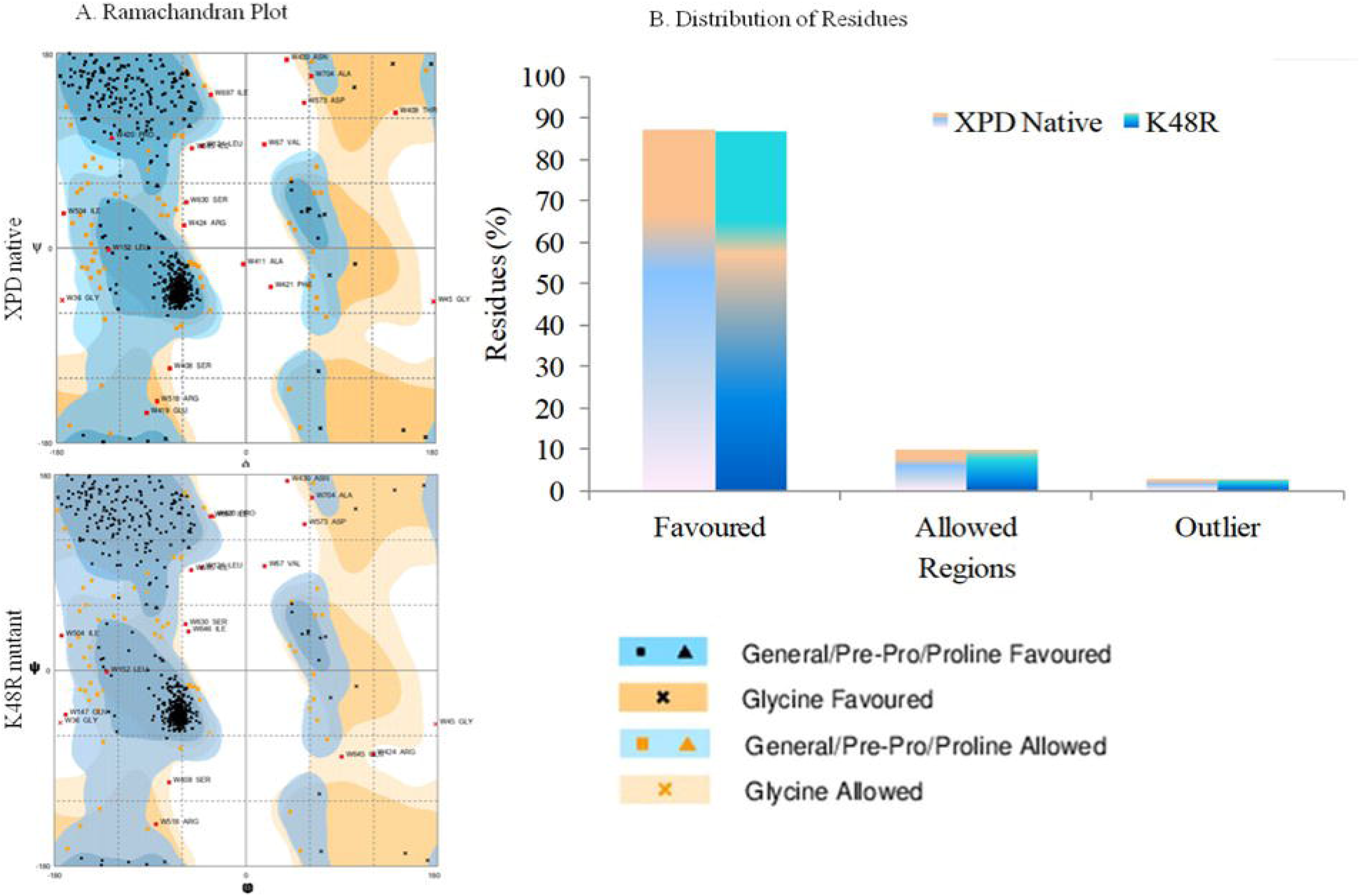
Quality Assessment of mutant and native models **A:** The Ramachandran plot analysis of mutant and native models **B:** Distribution of residues in Ramachandran plot.

### Analysis of Molecular Dynamics Simulations

To apprehend the conformational changes in the XPD owing to K48R mutation, molecular dynamics simulation (MDS) was performed for 25 nanoseconds. Several parameters have been analysed throughout the simulation trajectory including root mean square deviation (RMSD), root mean square fluctuations (RMSF), a total number of intramolecular hydrogen bonds, the radius of gyration, solvent accessible surface area (SASA) and secondary structure elements (SSE) of the protein with the time-dependent function of MDS.

RMSD profile for the backbone residue of the native and mutant protein was generated for all the atoms from the initial structure to understand the impact of the mutation on the stability of protein structure. [Figure 5A, 5B] shows that the RMSD value of the mutant structure is highly unstable as compared to the native protein. They showed a distinctly different type of deviation throughout the entire simulation, resulting in the backbone deviation range from ~0.4 nm (native) to ~1.0 nm (mutant) respectively. Native protein was found to be stabilized after 5 nanoseconds, whereas mutant protein lingered on till 15 nanoseconds to acquire stable conformation to a certain extent [Figure 5A, 5C].

**Figure 5.**
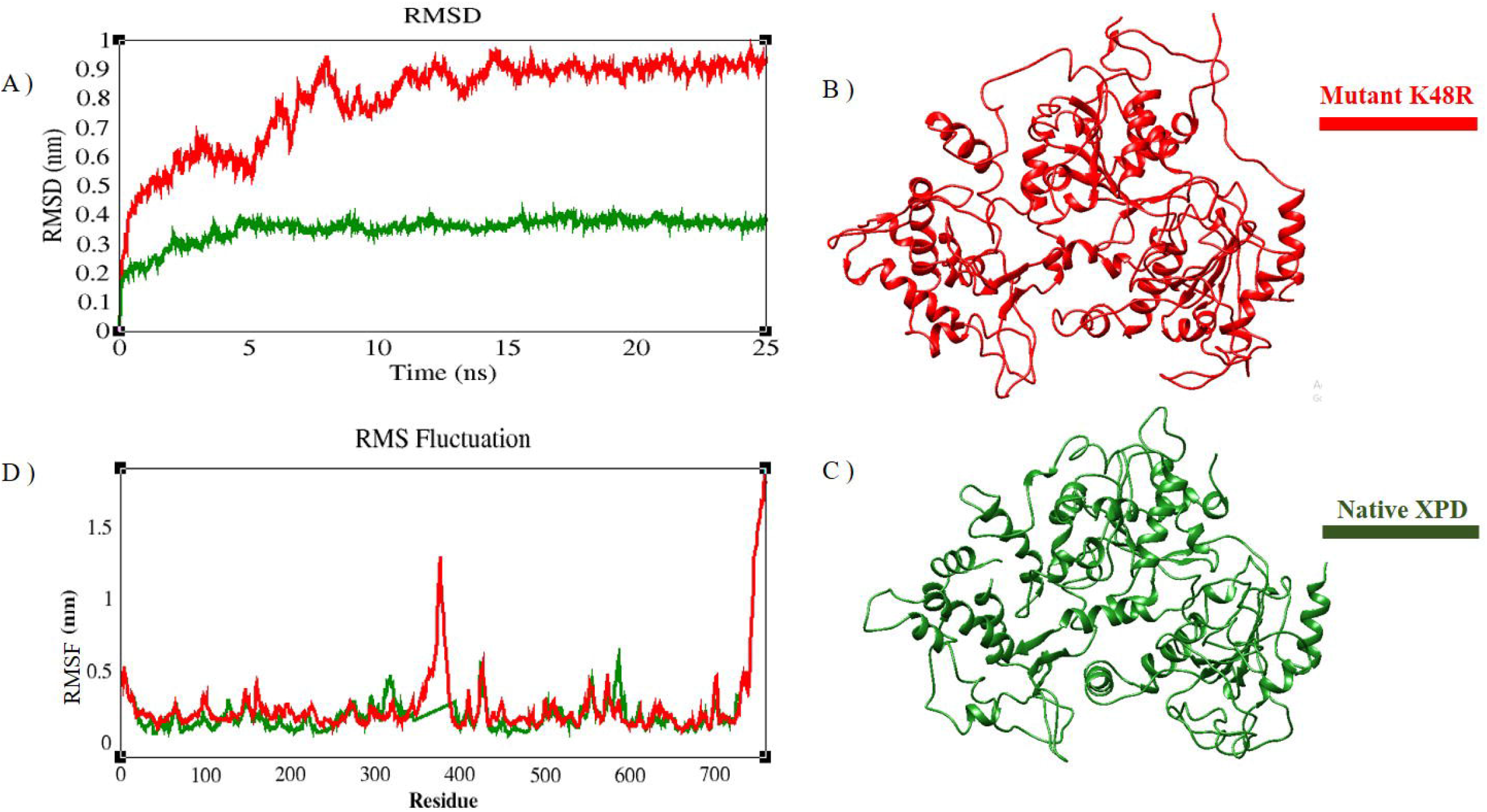
Molecular dynamics simulation of native and mutant model protein at 25000 ps. **A:** Time evolution of backbone RMSD is shown as a function of time of the native and mutant structures at 25000 ps. Native is shown in green and mutant in red. **B:** Simulated K48R mutant after MD. **C:** Simulated Native after MD **D:** RMSF of the backbone Cα atoms over the entire simulation. The ordinate is RMSF (nm) and the abscissa is an atom. Native is shown in green and mutant in red.

Next, we analysed how mutant affected the dynamic behavior of the residue through the calculation of RMSF of the Cα atom of the native and mutant protein. From [Figure 5D], it can be deduced that residue level fluctuations for mutant structure were rather high when compared with native structure, especially for residues located between 300 and 400 positions.

Subsequent intramolecular hydrogen bond analysis reaffirmed higher flexibility of mutant protein as we found a significant decrease in the number of hydrogen bond formation during the entire simulation as compared to the native structure [Figure 6C]. Native protein displayed the maximum number of hydrogen bonds in the range of 440 – 480 and whereas the number of hydrogen bonds drifted between 450 – 440 in the case of the mutant protein.

**Figure 6.**
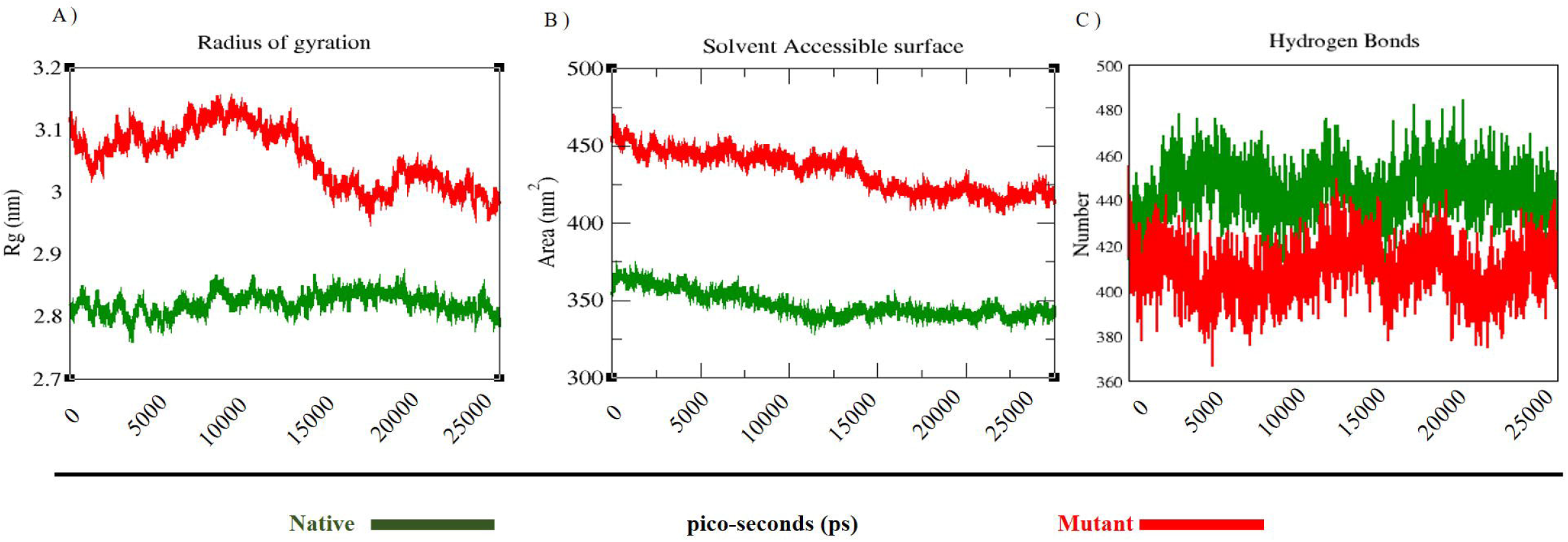
**A:** Radius of gyration of Cα atoms of native and mutant structure versus time. Native is shown in green and mutant in red. **B:** Solvent-accessible surface area (SASA) of native and mutant structure versus time. Native is shown in green and mutant in red. **C:** Average number of intermolecular hydrogen bonds in native and mutant structures as a function of time. Native is shown in green and mutant in red.

The radius of gyration (Rg) analysis revealed that mutant protein had the least compactness of its structure with 3.0 – 3.1 nm, while the native protein had shown to possess high compactness with 2.8 nm [Figure 6A].

The variation in SASA of mutant and native protein with time is depicted as [Figure 6B]. The native protein showed a lower SASA value in the range of ~340 nm^2^ – 360 nm^2^ and remained consistent throughout the simulation period whereas the mutant form exhibited higher value in the range of ~425 – 470 nm^2^ area with huge drift.

DSSP algorithm was applied to analyse the structural plasticity of protein. [Figure 7] showed the secondary structural elements as a function of simulation time. Coils, α-helix, turns, bends, and β-sheets were found in both native and mutant protein during the simulation time. However, the mutant protein showed a considerable decrease in α-helix forming residues with the concomitant increase in coils and bends forming residues as compared to native protein [Figure 7].

**Figure 7.**
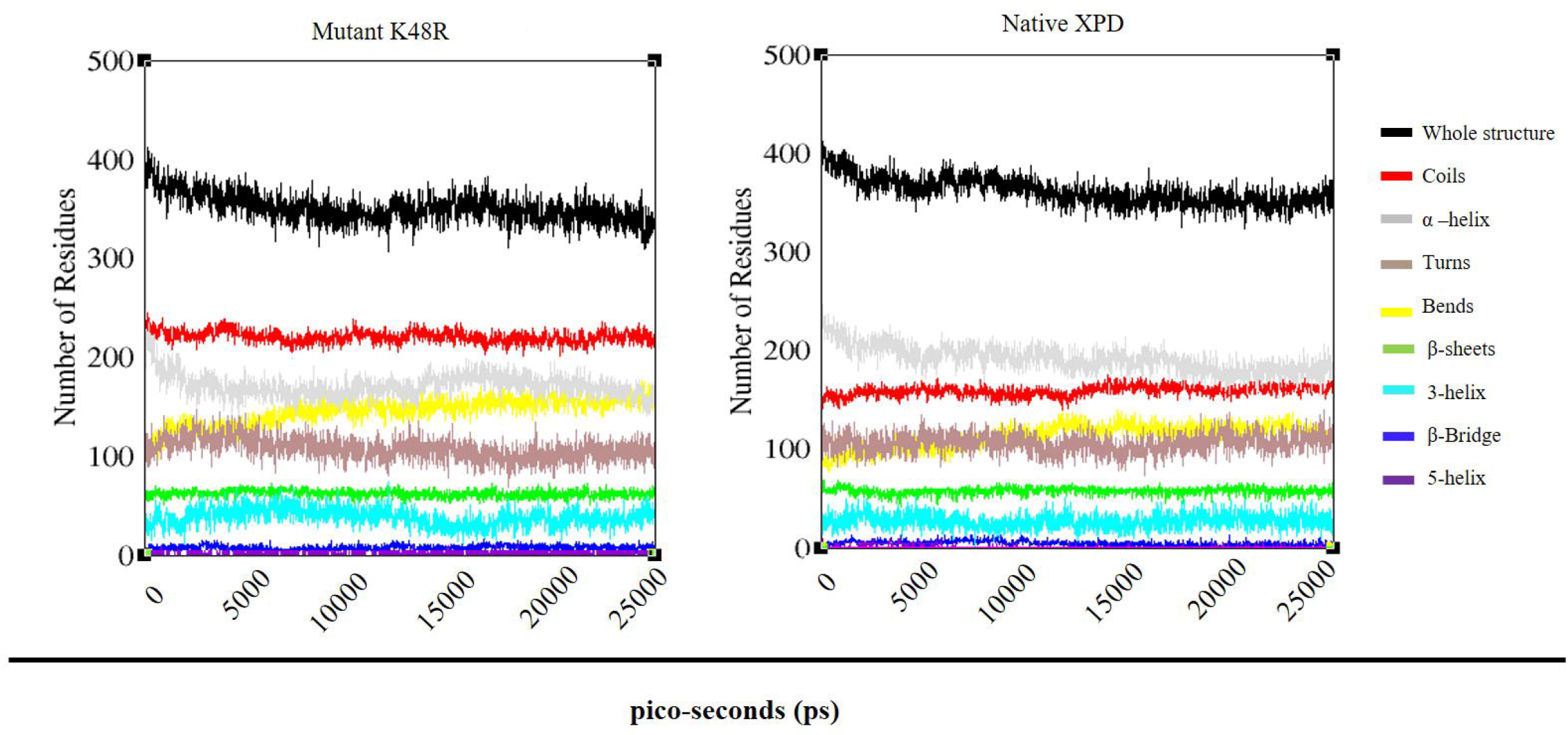
Time evolution of the secondary structural elements of a native and mutant structure at 25000 ps (DSSP classification)

Essential dynamics (ED) analysis was performed to identify the correlated motions of the native and mutant protein which gives the sum of the eigenvalues measured for total motility of the system. The spectrum of eigenvalues indicated that major fluctuations of the system were confined to the first two eigenvectors of both native and mutant. Hence the projection of trajectories of native and mutant simulations in the phase space along the first two principal components (PC1, PC2) at 300 K was plotted and depicted as [Figure 8]. Mutant protein covered a larger region of phase space along PC1 and PC2 as compared to the native protein.

**Figure 8.**
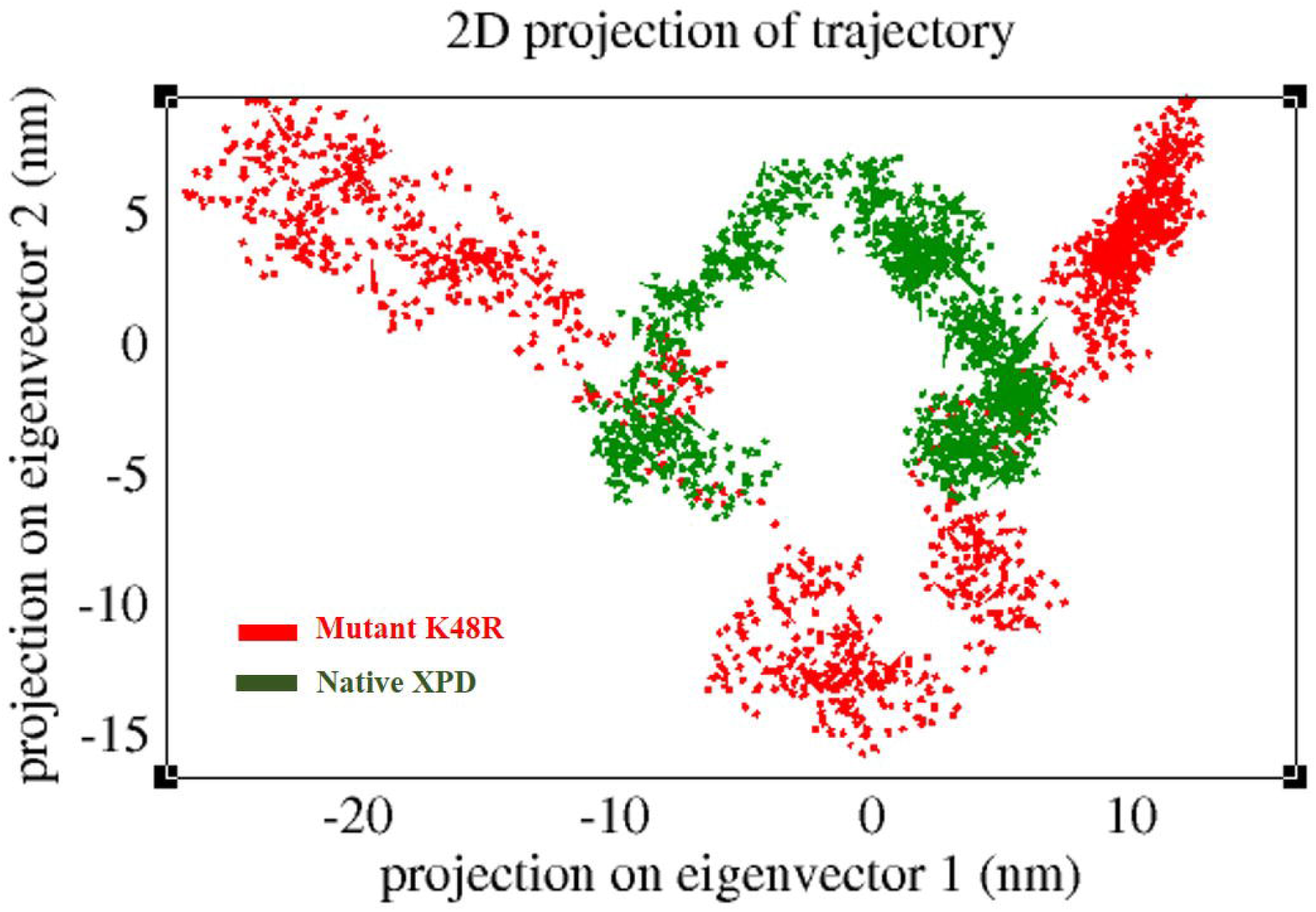
2D Projection of Principal Component Analysis. Projection of the motion of the protein in phase space along the first two principal eigenvectors. Native is shown in green and mutant in red.

## Discussion

XPD is a key NER enzyme that arbitrates in the transcription-coupled NER sub-pathway and can cause XP and other diseases when mutated in the germline [21]. However, recent advances in computational biology have afforded means to investigate genotype-phenotype correlation and their association with disease status [22–26]. Several bioinformatics algorithms used in combination frequently spotlight candidate functional missense mutations associated with genes [27–31]. In this study, we employed various bioinformatics tools to screen potentially deleterious cancer-causing mutations of *XPD* that could have a profound impact on protein stability and function.

Several computational algorithms predicted 64 mutations as deleterious out of 276 mutations retrieved from the public database. Bioinformatics tools predicting protein stability upon mutation showed varied outputs. However, FATHMM analysis predicted 7 mutations as cancer-causing/cancer-promoting mutations. Among the seven mutations, we selected K48R mutation and performed conservation analysis which revealed that the K48 position was conserved at a scale of 6. A previous study investigating the impact of K48R mutation on XPD activity showed that it inhibits both ATPase and helicase activities and eventually impedes the opening of the damaged DNA for potential repair process [21,32–34]. Interestingly, this mutation lies in the HD1 domain and mainly resides in the ATP binding pocket of XPD [21], making it an alluring candidate for further comprehensive investigations to understand its impact on protein structure, stability, and functionality.

Effect of mutations on protein structure and function can be envisioned by identifying the locus point of mutated amino acid within the protein structure. Several studies have reported the impact of relationships between mutation and their location in protein structure [9,27]. The molecular modeling approach using I-TASSER to model the mutant protein and subsequent MDS analysis facilitated us to investigate the structural effect of this mutation on protein structure. RMSD data imply that distinctly different type of deviation was noticed in the mutant protein throughout the entire simulation period as against native protein, thereby clearly demonstrating that mutation has a significant destabilizing effect on the protein. This was further corroborated by RMSF data which indicates that residue level variations were high for mutant protein, thereby indicating that mutation affected the protein conformation considerably leading to the increased flexibility of the protein. Subsequent intramolecular hydrogen bond analysis confirmed the higher flexibility of mutant protein as we found a significant decrease in the number of hydrogen bond formation during the entire simulation as compared to the native structure. Rg plot revealed the structural destabilizing effect on the protein owing to amino acid substitution at K48 that eventually leading to the loss of protein compactness. Besides, SASA analysis supported the Rg plot by returning higher values for mutant protein thus confirming the notion that protein might be enduring a major structural transition. Analysis of time-dependent secondary structure fluctuations through DSSP analysis showed a conformational drift from α-helix to coil form in mutant protein as the compared native protein. The conformational changes observed herein substantiate that major structural changes occurred due to amino acid substitution in the mutant structure which made it less stable, more flexible, and less compact. ED analysis demonstrated that the total motility of the mutant system was different from native protein and covered a larger region of phase space along PC1 and PC2 as compared to the native protein.

The association between the *in-silico* approach and wet lab experimentations has been rather evidenced by several previous studies [23,26]. Furthermore, computational mutation predictions when combined with MDS analysis has led to a breakthrough in the identification of most detrimental diseases causing mutations amid a huge pool of mutations [35]. Such computational approaches overlay the foundation for genetic studies to understand the molecular basis of disease and subsequent remedial measures [23,24] for future assessments of genetic variants in the clinic.

## Conclusion

Computational analysis of a large pool of missense mutations in the coding region of *XPD* revealed that substitution of lysine at 48^th^ residue with arginine could be deleterious mutation with potential cancer-causing/cancer-promoting role. MDS and ED analysis showed its altered structural stability owing to an increase in flexibility and a decrease in compactness. It is conceivable that loss of structural integrity in mutant protein might lead to loss of its interaction with ATP as well as with other partner proteins that in turn could affect its ability to unwind the damaged DNA double helix. This could effectively diminish DNA repair process that eventually could increase the risk for cancer. The mechanistic insights from this study may provide a cue for wet lab me to ascertain the impact of K48R mutation on cancer formation and design appropriate therapeutic strategy.

---

**Supplementary Table 1.**
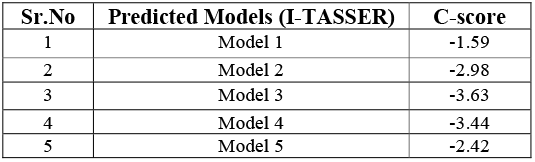
C-score predicted using I-TASSER Server

