## Supplementary Table for "Computational and Molecular Dynamics Simulation Approach To Analyze the Impact of *XPD* Gene Mutation on Protein Stability and Function"

**Supplementary Table 1:**

C-score predicted using I-TASSER Server

| **Sr.No** | **Predicted Models (I-TASSER)** | **C-score** |
| --- | --- | --- |
| 1 | Model 1 | -1.59 |
| 2 | Model 2 | -2.98 |
| 3 | Model 3 | -3.63 |
| 4 | Model 4 | -3.44 |
| 5 | Model 5 | -2.42 |
